# Contribution of *S1pr1*-featured astrocyte subpopulation to cisplatin-induced neuropathic pain

**DOI:** 10.1101/2025.04.03.642851

**Authors:** Ying Li, Silvia Squillace, Rachel Schafer, Luigino A Giancotti, Zhoumou Chen, Terrance M Egan, Stella G Hoft, Richard J DiPaolo, Daniela Salvemini

## Abstract

Chemotherapy-induced peripheral neuropathy accompanied by neuropathic pain (CIPN) is a major neurotoxicity of cisplatin, a platinum-based drug widely used for lung, ovarian, and testicular cancer treatment. CIPN causes drug discontinuation and severely impacts life quality with no FDA-approved interventions. We previously reported that platinum-based drugs increase levels of sphingosine 1-phosphate (S1P) in the spinal cord and drive CIPN through activating the S1P receptor subtype 1 (S1PR1). However, the mechanisms engaged downstream of S1PR1 remain poorly understood. Using single cell transcriptomics on male mouse spinal cord, our findings uncovered subpopulation-specific responses to cisplatin associated with CIPN. Particularly, cisplatin increased the proportion of astrocytes with high expression levels of *S1pr1* (*S1pr1*^high^ astrocytes), specific to which a Wnt signaling pathway was identified. To this end, several genes involved in Wnt signaling, such as the fibroblast growth factor receptor 3 gene (*Fgfr3*), were highly expressed in *S1pr1*^high^ astrocytes. The functional S1PR1 antagonist, ozanimod, prevented cisplatin-induced neuropathic pain and astrocytic upregulation of the Wnt signaling pathway genes. FGFR3 belongs to the FGF/FGFR family which often signals to activate Wnt signaling. Intrathecal injection of the FGFR3 antagonist, PD173074, prevented the development of CIPN in male mice. These data not only highlight FGFR3 as one of the astrocytic targets of S1PR1 but raise the possibility that S1PR1-induced engagement of Wnt signaling in *S1pr1*^high^ astrocytes may contribute to CIPN. Overall, our results provide a comprehensive mapping of cellular and molecular changes engaged in cisplatin-induced neuropathic pain and decipher novel S1PR1-based mechanisms of action.

## Introduction

The development of chemotherapy-induced peripheral neuropathy accompanied by neuropathic pain (CIPN) is a major dose-limiting neurotoxicity of chemotherapeutics [6] including cisplatin, a platinum-based chemotherapeutic widely used as a first-line therapy for lung, ovarian, and testicular cancer treatment [5]. In a clinical setting, cisplatin induces painful peripheral neuropathy with a prevalence up to 85% [10]. These symptoms can manifest years after chemotherapy completion, greatly impacting cancer survivors [58]. There are no FDA-approved therapies.

Molecular changes within the central nervous system (CNS) and the periphery contribute to the development of CIPN [3; 4; 13]. For example, in the periphery several chemotherapeutics including platinum-based drugs lead to partial degeneration of intraepidermal nerve fibers (IENFs), the afferent’s sensory terminal arbor, which may be responsible for the development of an abnormal incidence of spontaneous discharge of low frequency and irregular pattern in both A– and C-fiber afferents [3; 46]. The spontaneous discharge probably plays a role in the generation of allodynia and hyperalgesia [3]. Several studies including ours suggested that IENF degeneration is a consequence of chemotherapy-evoked mitochondria dysfunction, including loss of mitochondrial respiration and ATP production that result in a chronic energy deficiency in the peripheral sensory afferent [3].

On the other hand, in the spinal cord, alterations in neuro-immune communication are pivotal to the development and maintenance of neuropathic pain [15; 18; 43]. In this context, glial cells play important roles, releasing a variety of cytokines that increase neuronal excitability and perpetuate the extent of neuro-immune dysregulation by attracting immune cells that ultimately contribute to neuropathic pain [26; 27; 68; 70]. Over the years, we have shown that a linchpin in these processes is chemotherapy-induced activation of sphingolipid metabolism leading to increased production of the potent inflammatory sphingolipid, sphingosine-1-phosphate (S1P) and S1P receptor subtype 1 (S1PR1) activation in the spinal cord [19; 61; 63]. Accordingly, inhibition of S1PR1 with functional and competitive S1PR1 antagonists blocks and reverses CIPN [19; 63; 64]. These findings are noteworthy, in that several functional S1PR1 antagonists such as fingolimod, ponesimod and ozanimod have been FDA-approved for treatment of relapsing multiple sclerosis (MS) [25; 53; 56] and more recently for ulcerative colitis [2]. Moreover, S1PR1 antagonists do not interfere with chemotherapy’s anticancer activity [24; 33; 40; 57] and are being considered as novel anticancer agents [52; 57]. Therefore, functional S1PR1 antagonists have the potential to be repurposed as adjuncts to chemotherapy to mitigate major neurotoxicity [61]. As recently reported by our group, spinal cord astrocytes are an important cellular locus of S1PR1 activity; thus, deletion of S1RP1 in astrocytes attenuated the pharmacological activity of S1PR1 antagonists [63].

However, the molecular mechanisms involved remain obscure. Our work fills gaps in our understanding of the roles of S1P in cisplatin-induced neuropathic pain by unravelling previously unrecognized links between S1PR1 and Wnt signaling in spinal cord astrocytes. Our work also provides the first evidence of novel neuroprotective effects of functional S1PR1 antagonism in the periphery through prevention of IENF loss continuing to support repurposing S1PR1 antagonists as adjuncts to chemotherapy.

## Methods

### Animals

Procedures for the maintenance and use of animals were in accordance with the NIH Guide for the Care and Use of Laboratory Animals (National Academies Press, 2011) and approved by the Saint Louis University Institutional Animal Care and Use Committee. Seven to eight week-old male wildtype (WT; C57BL/6) mice were purchase from Envigo. All mice were housed 4-6/cage in a controlled environment (12-h light-dark cycles) with food and water available *ad libitum*. All experiments were conducted with the experimenters blinded to treatment conditions.

### Cisplatin-induced neuropathic pain model

To induce neuropathic pain, mice received two cycles of cisplatin as previously described [9]. Briefly, mice received daily intraperitoneal (i.p.) injections of cisplatin (2.3 mg/kg, Accord Healthcare, Inc., Durham, NC, USA) or saline for 5 consecutive days, then rested (no injections) for the following 5 days followed by another 5-day cycle of cisplatin or saline. The cumulative dose of cisplatin used in this study (23 mg/kg, human equivalent 70 mg/m^2^) is comparable to the total dose per cycle used in patients (75 mg/m^2^). Body weight and overall well-being of the animals were monitored throughout the treatment period.

When testing candidate compounds using intrathecal (i.th.) injection, mice were given daily injections of cisplatin (2.3 mg/kg, i.p.) or saline for a cycle of 5 days only.

### Tested compounds preparation and administration

Ozanimod was purchased from Med Chem Express (Monmouth Junction, NJ, USA). The dose of ozanimod used in this study (1mg/kg, in 5% DMSO and 5% Tween 20 in saline) was chosen based on previous studies from our group that demonstrated it was the optimal dose to completely block S1PR1-mediated neurotoxicities [8; 19; 60; 63]. Mice received i.p. injections of ozanimod (1 mg/kg in 5% DMSO and 5% Tween 20 in saline) or their vehicle 15 minutes before each cisplatin injection.

FGFR3 antagonist, PD173074, was purchased from Med Chem Express (Monmouth Junction, NJ, USA). Mice received i.p. injections of PD173074 (20 mg/kg in 5% DMSO and 5% Tween 20 in saline) or their vehicle 15 minutes before each cisplatin injection. For repeated intrathecal injection, a total of 10 µl PD173074 (5 µM in 5% DMSO and 5% Tween 20 in saline) or vehicle was injected 15 minutes before each cisplatin injection.

### Behavioral Testing

Mechano-hypersensitivity as a readout for neuropathic pain development was assessed through the hind paw withdrawal response to von Frey hairs stimulation, according to the up-and-down method [11]. In brief, animals were acclimatized (30 minutes-1 hour) to individual clear Plexiglas boxes on an elevated wire mesh platform before probing with von Frey filaments beginning with 0.4 g bending force. Filament bending force was increased until there was a positive response, defined by clear paw withdrawal or shaking. After the first positive response, the paw was probed with filaments of decreasing force until a negative response was observed and then with filaments of increasing force until a positive response occurred again. At least 4 readings were obtained after the first positive response, and the pattern of response was converted to a 50% paw withdrawal threshold (PWT) using the method described by Chaplan [7]. If a mouse did not respond at 2.00 g, they were given a positive response at 2.00 g. Animals exhibiting baseline PWT below 0.6 g were excluded from this study.

### Tissue dissociation and single cell isolation

Saline-perfused spinal cord lumbar segments (L4-L6, n=5/group) were harvested from mice 1h following behavioral measurements. Samples of each group were pooled and transported on ice to a Biosafety level-2 cabinet as soon as possible following resection, weighed, and then immediately submerged in 50 ml of ice-cold divalent-free Hank’s balance salt solution (HBSS) for 10 min. The cooled tissue was transferred to a 10 cm petri dish containing just enough cold HBSS to cover the tissue. The tissue was chopped into small pieces (∼1 mm^3^) using a sterile single-edge razor blade. Tissue chunks were enzymatically dissociated using papain and DNase. In brief, the minced tissue was incubated under slow continuous rotation at 37°C in HBSS containing papain (10 ml of 2 U/ml) for 10 min, mechanically agitated using a fire-polished Pasteur pipette, then incubated with rotation in a mix of papain and DNase for an additional 10 min. After that, leupeptin was added to make a final concentration of 20 µM and the enzymes were removed by centrifugation at 300 x g for 10 minutes. Then the tissue was washed with 50 ml of cold HBSS, and single cells were separated by passage through the ends of three to four Pasteur pipettes with fire-polished tips of diminishing diameters.

The resulting cell suspension was passed through a 40-µm cell strainer and the flow-through collected in a 50 ml falcon tube. The flow-through was centrifuged at 300 x g for 10 min at room temperature. The supernatant was removed. Myelin in the remaining pellet was removed by resuspending in 10 ml of room temperature 22% Percoll in PBS and overlaying with 5 ml PBS. The gradient was centrifuged at room temperature for 20 min at 950 x g with slow acceleration and no brake. The supernatant including the myelin interface layer was carefully removed and discarded. The remaining pellet was washed with 50 ml of HBSS and centrifuged at 300 x g for 10 min.

### Single cell RNA sequencing

Droplet-based single cell RNA-seq libraries were prepared with the 10x Genomics Chromium Controller using the Chromium Next GEM Single Cell 3 Reagent Kits v3.1 (10x Genomics, Pleasanton, USA), following manufacturer protocol. Each library was generated to have a maximum recovery of 10,000 cells. Polymerase chain reactions were conducted in the T100 Thermal Cycler (Bio-Rad, Hercules, CA). Final quality control was performed on the 2100 Bioanalyzer Instrument (Agilent, Santa Clara, CA), and libraries were sequenced on the NovaSeq X Plus (Illumina, San Diego, CA) at the Genome Technology Access Center of Washington University in St. Louis. Raw sequence reads were aligned to the mm10 (Ensembl 84) reference genome, using CellRanger software (v5.0.1). We uploaded the CellRanger output to Zenodo as Sample_1_filtered _feature_ bc_matrix, Sample_2_filtered_feature_bc_matrix, Sample_3_filtered_feature_bc_matrix, and Sample_4_filtered_feature_bc_ matrix, which can be accessed on Zenodo using DOI <u>10.5281/zenodo.14961645</u>.

### Clustering analysis

R package Seurat (v5.0.1) was used to generate cell clusters based on transcriptional profile. After eliminating low-quality cells, cells with >250 genes, > 500 transcripts, and < 20% mitochondrial transcripts were retained. Genes expressed in less than 10 cells were excluded from downstream analysis. Briefly, RNA counts were normalized with the LogNormalize method, and the top 3000 variable genes were extracted with the FindVariableFeatures function. Individual datasets were combined using IntegrateData and scaled using SCTransform function by regressing for mitoratio. PCA was performed on the 3000 variable genes and the top 36 principal components were selected for clustering with a resolution of 0.3. The clustering results were visualized using UMAP plots. The function FindConservedMarkers was used to find the DEGs/feature genes in each cluster with cutoff of min.diff.pct = 0.25, min.pct = 0.25, and logfc.threshold = 0.25. Cell types were identified by using several marker genes from the literature [28; 37].

For astrocytes subtype analysis, astrocytes were subset from the complete dataset and then rescaled using SCTransform function by regressing for mitoratio. PCA was performed on the 3000 variable genes and top 40 principal components were selected for clustering. The function FindConservedMarkers was used to find the DEGs/feature genes in each cluster with cutoff of min.diff.pct = 0.01, min.pct = 0.1, and logfc.threshold = 0.25.

### Differentially expressed genes identification and pathway enrichment analysis

For each cell type, DEGs across different sample groups were identified using the FindMarkers function and bimod test, with cutoff of min.pct = 0.1, and logfc.threshold = 0.25. Upregulated genes were defined with p_val < 0.05 & logfc.threshold > 0.25 and downregulated genes were defined with p_val < 0.05 & logfc.threshold < – 0.25. Pathway enrichment analysis was performed on the DEGs using clusterProfiler function in R [74].

### RNAscope

<u>Tissue preparation.</u> Twenty-four hours after the last injection, animals were perfused with 1X PBS then 4% PFA in 1X PBS sequentially. The lumbar segment of spinal cords was dissected and post fixed overnight in 4% PFA in 4 °C. The tissues were then sequentially treated for cryopreservation in 20% and 30% sucrose in 4 °C for 24h each. The fixed and cryopreserved lumbar segment were snap frozen in liquid nitrogen cooled 2-methyl butane.

Fixed frozen tissues were sliced into 7 µm sections on glass slides using cryostat and then stored at –80°C until the experiment date. Stained sections were prepared according to the manufacturer’s instructions (Advanced Cell Diagnostics, Newark, CA). Briefly, tissue sections were dried at 60°C for 30 minutes before treating with hydrogen peroxide, antigen retrieval and protease. A cocktail of probes for *Fgfr3* (Mm-Fgfr3), *S1pr1* (Mm-S1pr1-C2) with *Gfap* (Mm-Gfap-C3) was incubated for 2h at 40°C in a humidified chamber before amplification and detection with TSA Vivid Fluorophore 520 (PN 323271), TSA Vivid Fluorophore 570 (PN 323272) and TSA Vivid Fluorophore 650 (PN323273). (Advanced Cell Diagnostics, Newark, CA). Slides were counterstained with DAPI and mounted in ProLong® Gold (ThermoFisher Scientific) and imaged using Andor Dragonfly Spinning Disk Confocal. For quantification, 20x image of the dorsal horn were scanned at 2048 x 2048 pixels. QuPath was used to measure the count of *Fgfr3* signal/dots that were associated with *Gfap* or *S1pr1* [55].

<u>Image analysis.</u> Images were imported to QuPath software. Basically, a ROI was selected to cover the whole lamina I, II, and III of the dorsal horn based on DAPI signals of each animal. Thresholds of examined signals were set identical through images for analysis. Number of mRNA signals at single cell level was averaged and then graphed using Prism (version 10.4.1 for Windows, GraphPad Software, San Diego, CA).

### Immunofluorescence

When testing FGFR3 and GFAP protein levels in DH-SC, glass slides containing 7 µm sections of fixed frozen spinal cord (tissue preparation described above) were used. Glass slides with tissue sections were blocked for 1h at room temperature in blocking buffer (10% donkey serum in 1X PBS + 0.25% Tween-20 + 1% BSA). The slides were then washed in 1x PBS for 3 times, 5 minutes each and incubated with primary antibodies to GFAP (1:1000; #PA5-18598; ThermoFisher Scientific), and FGFR3 (1:100; # PA5-34574; ThermoFisher Scientific) in blocking buffer at 4 °C degree overnight. Slides were washed extensively and stained for 1h at room temperature with a cocktail (1:300 dilution each) of anti-goat Alexa 488 (#705-545-147; Jackson Immunoresearch), anti-rabbit Alexa 594 (#711-585-152; Jackson Immunoresearch), and DAPI (1:2500, ThermoFisher). Sections were mounted in Prolong® Gold (#P36930; ThermoFisher) and imaged using Andor Dragonfly Spinning Disk Confocal. For quantification, 20x image of the dorsal horn were scanned at 2048 x 2048 pixels.

<u>Image analysis.</u> Images were imported to Huygens Professional and analyzed by Huygens Object Analyzer (SVI, Netherlands). Basically, a ROI was selected to cover the whole lamina I, II, and III of the dorsal horn based on DAPI signals of each animal. Thresholds of anti-GFAP and anti-FGFR3 signals were set identically through images for analysis. Immunofluorescence intensity of antibody signals was divided by the corresponding ROI area and then graphed using Prism (version 10.4.1 for Windows, GraphPad Software, San Diego, CA).

For IENFs staining, twenty-four hours after the last injection, mice were deeply anesthetized with isoflurane and perfused with PBS, followed by 4% paraformaldehyde (PFA). The hind paw glabrous skin was collected, post fixed for 24 hours in 4% PFA, cryoprotected in 20% sucrose overnight and then 30% sucrose overnight, and then embedded in optimal cutting temperature (OCT) compound (Sakura Finetek). Tissue was sectioned with a Cryostat (Leica CM1950) at a thickness of 15 μm and thaw mounted onto SuperFrost Plus slides (Thermo Fisher Scientific) and slides were stored at –80°C until use. Prior to use, slides were thawed for at least 30 minutes at room temperature and washed in PBS to remove excess OCT. Slides were blocked with PBS supplemented with 0.3% Triton X-100 and 3% normal donkey serum. Slides were incubated with primary antibodies (rabbit anti-human PGP9.5, CL7756AP, 1:1000) diluted in PBS supplemented with 0.1% Triton X-100 and 1% normal goat or donkey serum at 4°C degree overnight. Slides were washed and incubated with anti-rabbit Alexa 594 (#711-585-152; Jackson Immunoresearch), and DAPI (1:2500, ThermoFisher) at room temperature for 1 hour. Slides were washed, then coverslipped with Prolong Gold mounting media (Thermo Fisher Scientific). Images were captured on a Leica SP8 Laser Confocal microscope with objective of NA = 1.30, 40x. The number of IENFs in each image were calculated by NIH Image software and divided by the length of basement membrane. Values were then graphed by Prism (version 10.4.1 for Windows, GraphPad Software, San Diego, CA).

### Statistics

All data are expressed as mean ± SD of n number of animals and analyzed by GraphPad Prism (version 10.4.1 for Windows, GraphPad Software, San Diego, CA). Data for log10-transformed von Frey behavioral studies were analyzed by Two-way ANOVA with Tukey’s multiple comparisons test. Immunofluorescence intensity data and RNAscope probes signal quantification data were expressed as mean ± SD and analyzed by one-way ANOVA with Tukey’s multiple comparisons test. A P < 0.05 was considered significant difference for behavioral and biochemical studies.

## Results

### Ozanimod prevents cisplatin-induced neuropathic pain

CIPN was induced as previously described [60]. Briefly, mice were injected with cisplatin intraperitoneally (i.p.; 2.3 mg/kg) for two cycles of 5 daily injections with 5 days of rest in between [60]. Cisplatin caused significant mechano-allodynia, which was prevented by ozanimod administered intraperitoneally (i.p.; 1 mg/kg) 15 minutes before each cisplatin injection (**Fig. 1A**).

**Figure 1.**
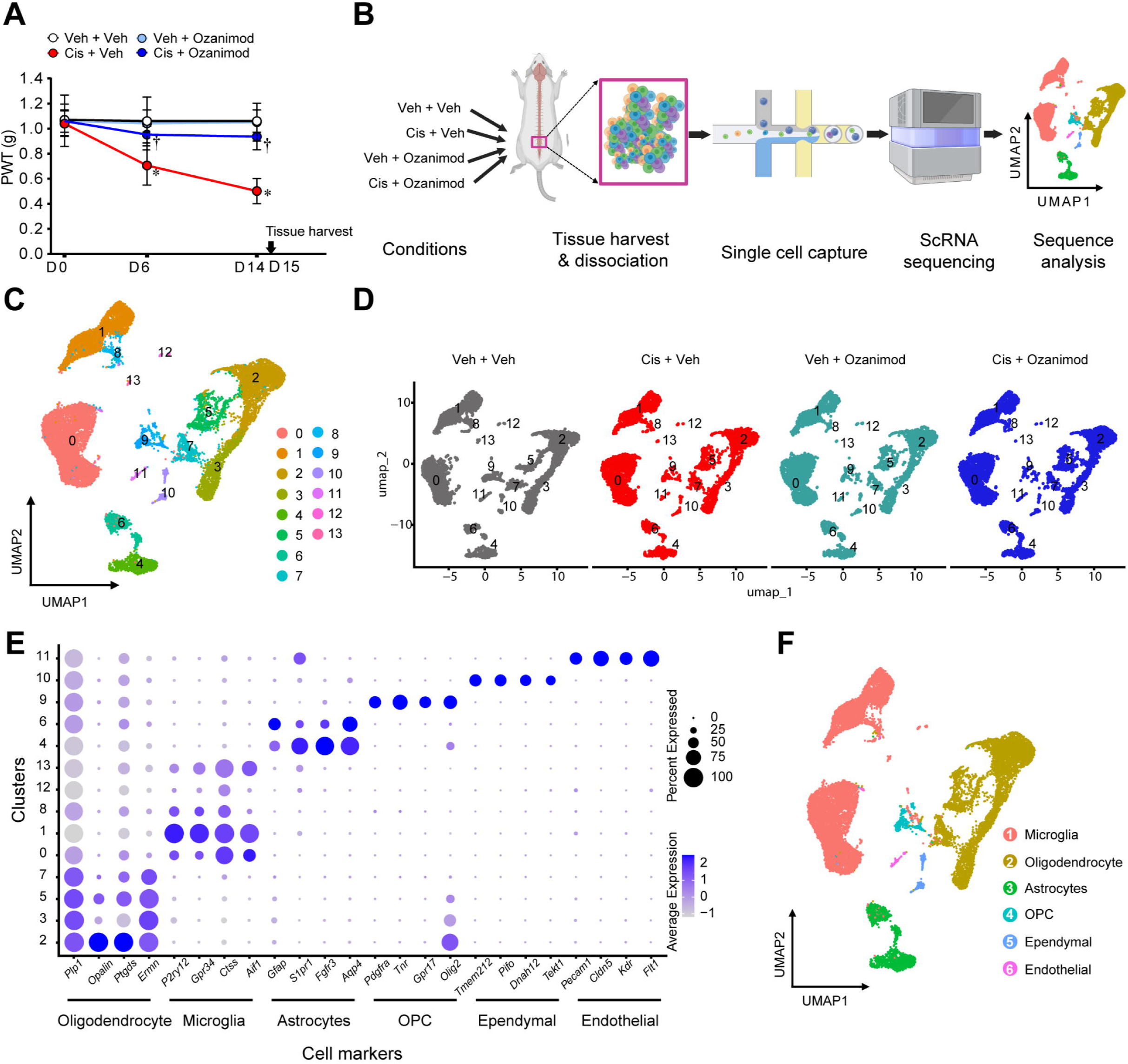
Workflow of scRNA-seq of spinal cord from mice with cisplatin-induced neuropathic pain (CIPN). (A) Mechanical hypersensitivity assessment of mice used for single-cell RNA sequencing. Mechanical hypersensitivity was assessed before the start of the treatment (D0), 48h after the end of the first cycle of cisplatin (D6) and on the day of the last treatment (D14). Mean±SD, n=5/group, Two-way ANOVA with Tukey’s multiple comparisons test. *P<0.05 vs. Veh + Veh, † P<0.05 vs. Cis + Veh. Twenty-four hours after the last treatment, spinal cord was harvested for single-cell RNA sequencing. (B) Workflow of scRNA-seq. The lumbar spinal cord was subjected to single cell isolation followed by sequencing and data analysis. (C) UMAP plot of 18516 cells from all conditions. Cells were present in 21 distinct clusters. (D) UMAP plot of cells from each condition. (E) UMAP plot and violin plot (F) of expression pattern of marker genes that identify each cell type. The values indicated are log2 normalized counts per cell. (G) UMAP plot with color-coded cell types. OPC, oligodendrocyte precursor cells.

### Single cell RNA (ScRNA)-seq analysis of spinal cord tissues

To assess the cellular heterogeneity and identify downstream targets of S1PR1, we performed single cell RNA sequencing on spinal cord tissues harvested 15 days after the first vehicle, ozanimod, cisplatin or cisplatin plus ozanimod treatments as described in **Fig. 1A,B**. After quality control and potential doublets filtration, we obtained a total of 18516 cells from mice treated with vehicle (Veh + Veh, 5784 cells), cisplatin alone (Cis + Veh, 4189 cells), ozanimod alone (Veh + Ozanimod, 3906 cells), and cisplatin with ozanimod pretreatment (Cis + Ozanimod, 4637 cells). Cluster analysis on the cells from all samples revealed 14 distinct clusters, which contained comparable cells from each sample (**Fig. 1C,D**). Cluster-specific marker genes were identified by comparing the expression of each gene in one cluster to the average of the others (**Supplemental Fig. 1A**). Clusters were annotated when established cell-specific marker genes were detected (**Fig. 1E**). Overall, we identified 6 major cell types: microglia, oligodendrocyte, astrocytes, oligodendrocyte precursor cells (OPC), ependymal, and endothelial, with their proportions in total cells in descending order (**Fig. 1F**). Neurons were not expected to survive our single-cell dissociation protocol and consequently were not identified in the analysis. Microglia, oligodendrocyte, and astrocytes together took up more than 90% of total cells; cisplatin induced significant changes (∼10% of increase for microglia and ∼10% of decrease for oligodendrocyte and astrocytes) in their cell-type proportions (**Supplemental Fig. 1B**). Notably, these changes were not observed in sample group receiving both ozanimod and cisplatin, indicating a protective effect of ozanimod on cisplatin-induced cell-type composition alterations (**Supplemental Fig. 1B**).

### Cisplatin induces transcriptional reprogramming in spinal cells

Pairwise comparisons of gene expression between the vehicle and cisplatin alone group at cell-type level resulted in different numbers of differentially expressed genes (DEGs), ranging from 844 in microglia to 4370 in OPC. Pathway enrichment analysis was performed on these DEGs to identify cisplatin-induced dysregulations in cellular processes. For each cell type, we mapped the top 50 upregulated or downregulated pathways and picked 5 representative ones that have no overlaps in the associated DEGs, with an example shown in **Supplemental Fig. 2**. We found that cisplatin consistently downregulated the heat shock protein 90 alpha family class A gene *Hsp90aa1* and *Hsp90ab1* in all cell types (**Fig. 2A**), resulting in a suppression effect on protein folding and stability in CIPN mice (**Fig. 2B**). Pathway analysis also uncovered cell type-specific dysregulation in response to cisplatin. For instance, cisplatin downregulated astrocytic ATP synthesis-coupled electron transport, which is essential for ATP-mediated cell-to-cell communications in CNS [20]. Cisplatin also specifically upregulated the ERK1 and ERK2 cascade pathway in microglia (**Fig. 2B**), which is known to function in activating inflammatory responses to drive neuropathic pain [65]. Overall, these results indicate that cisplatin suppressed protein folding and stability in all the identified cell types while it also induced cell type-specific dysregulation that may contribute to the development of CIPN.

**Figure 2.**
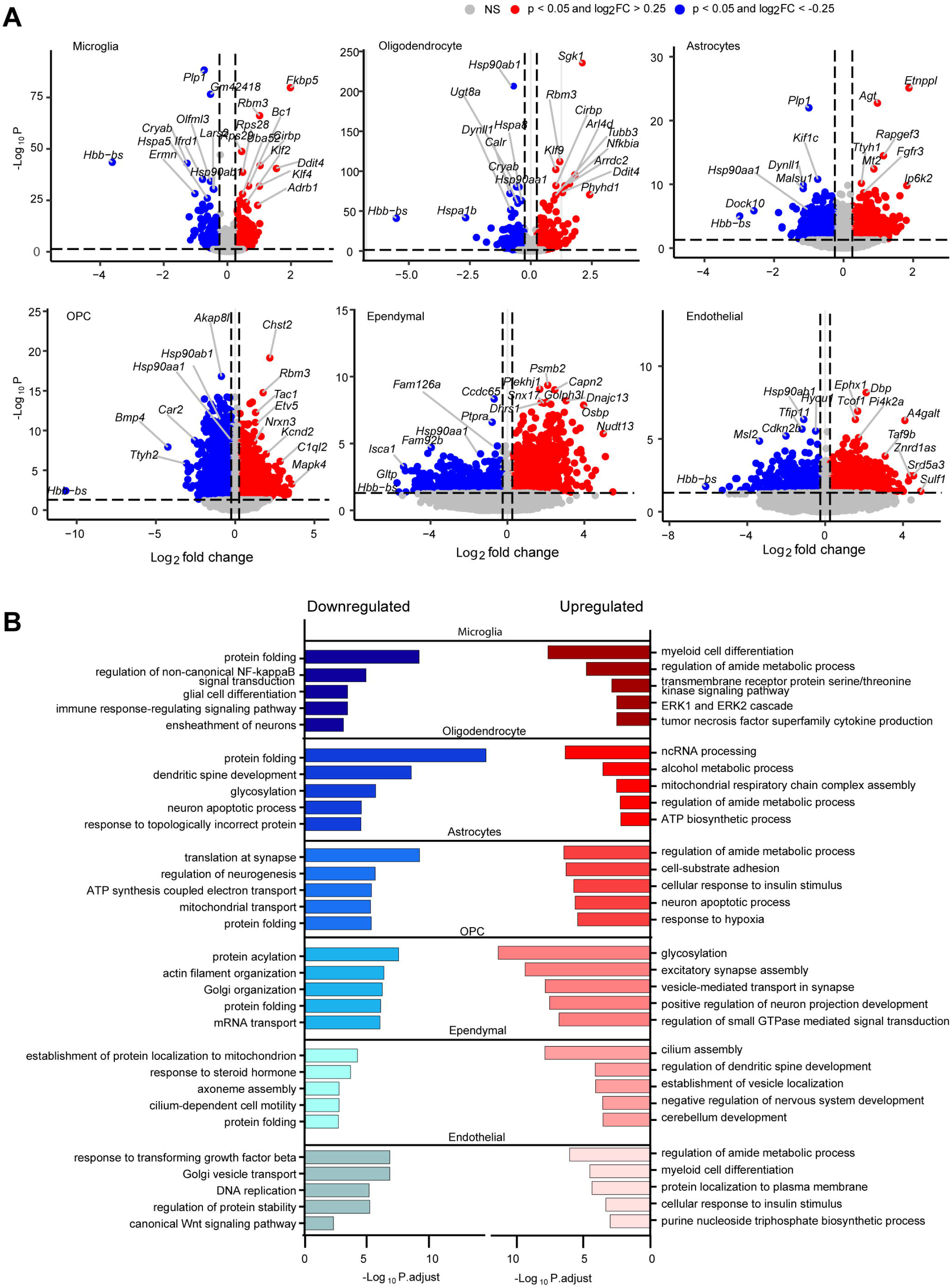
Cisplatin effects on transcriptional profile of spinal cells. (A) Volcano plot of upregulated (red dots) and downregulated genes (blue dots) in different cell types. DEGs with p-value (p-val) < 0.05 and absolute Log2 fold change > 0.25 were defined as significant regulated genes. (B) Pathway analysis of upregulated and downregulated genes shown in (A), using cutoff of adjusted p-value (P.adjust) < 0.05 and a minimum overlap of three genes in the pathway.

The effects of cisplatin in spinal neurons were not determined due to the absence of neurons in the study. However, neuronal damage is inferred by the observed cisplatin-induced dysregulation of neuronal processes in microglia and astrocytes. Particularly, cisplatin downregulated the neuron ensheathment pathway in microglia and induced the neuron apoptotic process and suppressed neurogenesis regulation in astrocytes (**Fig. 2B**). These results are consistent with the protective role of glia cells in neuronal development [16; 39; 54] and implicating the contribution of cisplatin-induced glial transcriptional reprogramming to potential neuron damage in CIPN mice.

### Astrocyte heterogeneity analysis reveals *S1pr1*-featured subpopulation

*S1pr1* is highly expressed in astrocytes. Mice with astrocyte-specific deletions of *S1pr1* do not develop mechano-hypersensitivity in multiple neuropathic pain models, including CIPN [8; 63]. To better understand astrocyte heterogeneity and characterize *S1pr1*-expressed astrocytes in the context of CIPN, we performed further cluster analysis on astrocytes and identified 3 subpopulations, cluster 1, cluster 2, and cluster 3 (**Fig. 3A**). *Gfap* encodes glial fibrillary acidic protein (GFAP) and is widely used as a cell marker gene for astrocytes [59]. Notably, *S1pr1* and *Gfap* were highly expressed in cluster1 and cluster2, respectively (**Fig. 3B**). We used RNAscope *in situ* hybridization (ISH) to validate the distinct expression profiles of *S1pr1* and *Gfap* in spinal cord. Co-staining of *S1pr1*, *Gfap,* and *Fgfr3,* another cluster1 signature gene (**Fig. 3B**), in the dorsal horn of the spinal cord (DH-SC) showed *Gfap* mRNA signals mostly localized to the white matter and lamina I,II, and III of the grey matter, while *S1pr1* and *Fgfr3* mRNA signals were distributed all over the grey matter (**Supplemental Fig. 3A-D**). Detection of both *S1pr1* mRNA and *Gfap* mRNA signals at the single-cell level indicated their co-expression in astrocytes (**Supplemental Fig. 3E**). However, linear regression analysis of the number of *S1pr1* mRNA and *Gfap* mRNA signals in individual cells resulted in an R-squared value of 0.2270, suggesting that their expression levels were not correlated (**Supplemental Fig. 3H**). Similar results were obtained from linear regression analysis between mRNA signals of *Gfap* and *Fgfr3* (**Supplemental Fig. 3F,I**). On the other hand, analysis of the number of *S1pr1* mRNA and *Fgfr3* mRNA signals resulted in a linear regression with an R-squared value of 0.7061, indicating that expression of *S1pr1* and *Fgfr3* were highly correlated (**Supplemental Fig. 3G,J**), further supporting our scRNA-seq data.

**Figure 3.**
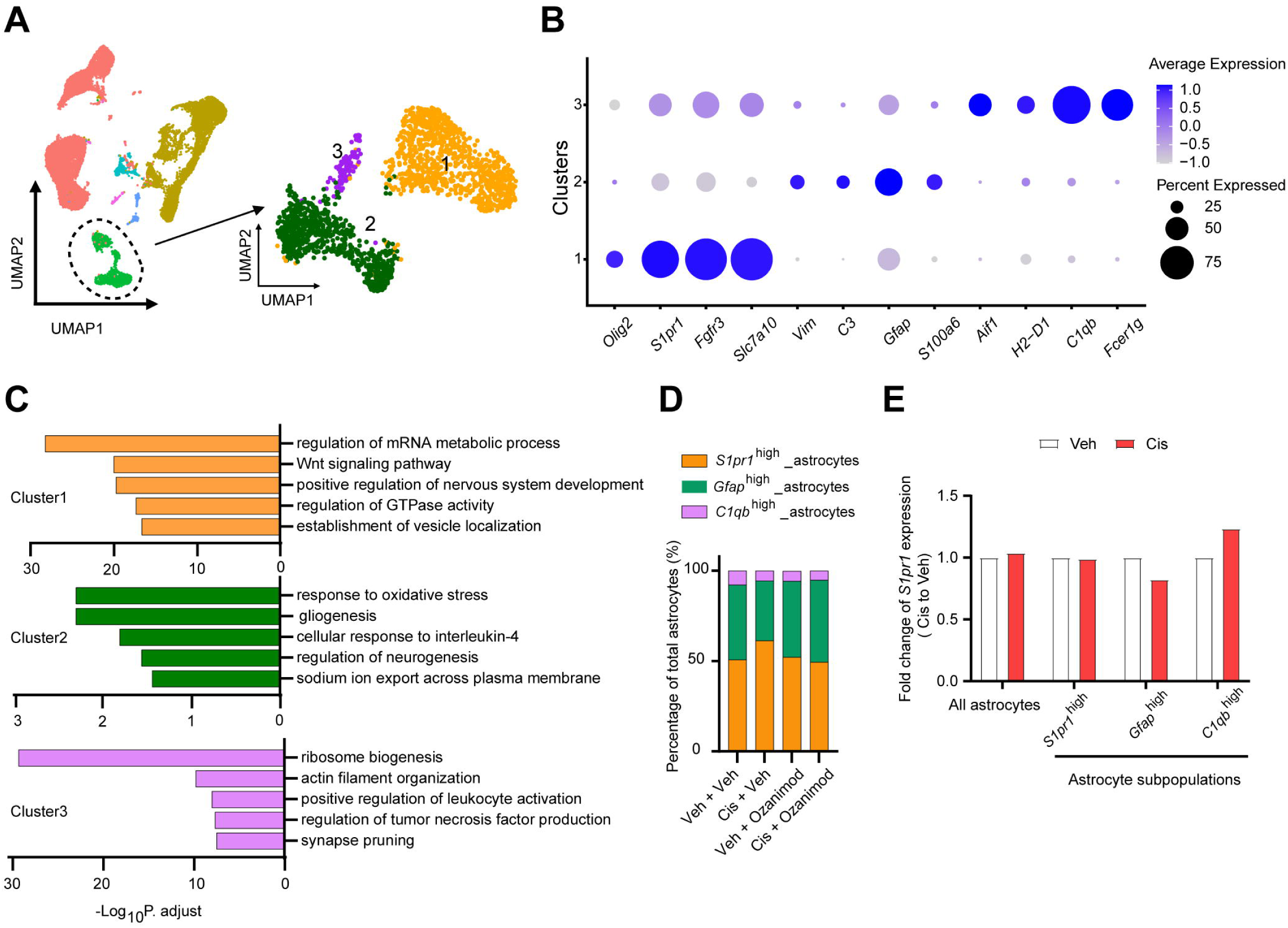
Molecular profile of astrocytes heterogeneity in mouse spinal cells. (A) UMAP plot showing 3 astrocytic clusters (subpopulations). (B) Dotplot of signature genes for each astrocyte cluster. (C) Pathway analysis of cluster-specific genes. Five representative pathways were picked from the top 50 enriched pathways. (D) Stacked bar chart showing proportion of each astrocyte subpopulation among all sample groups. (E) Effects of cisplatin on mean *S1pr1* expression levels in all astrocytes and individual subpopulations.

Pathway enrichment analysis on signature genes revealed cluster-specific biological processes. Cluster 1 astrocytes (*S1pr1*^high^) were associated with mRNA metabolic processes, the Wnt signaling pathway and GTPase activity regulation, in addition to the neuronal functions including nervous system development regulation and vesicle localization establishment (**Fig. 3C**). The Wnt signaling pathway was comprised of signature132 genes that not only have classic Wnt signaling components such as the Wnt ligand gene *Wnt7b*, frizzled receptors genes, *Fzd1*, *Fzd2*, and *Fzd10*, and adenomatous polyposis coli gene (*APC*) [31], but also have Wnt signaling regulator genes such as the FGF/FGFR family genes *Fgfr2* and *Fgfr3* [14; 17; 22]. Cluster 2 astrocytes (*Gfap*^high^) were associated with oxidative stress and interleukin-4 responses, gliogenesis and neurogenesis, as well as sodium ion export (**Fig. 3C**). Interestingly, astrocytes of cluster 3 featured high expression levels of microglia-specific genes such as *C1qb* and *Aif1* (**Fig. 3B**). Thus, the cluster 3 (*C1qb*^high^) astrocytes were associated with cytokine production and immune cell activation usually seen in microglia (**Fig. 3C**).

The *S1pr1*^high^ subpopulation represented the largest part of astrocytes, followed by *Gfap*^high^ and *C1qb*^high^ (**Fig. 3D**). Cisplatin treatment increased the proportion of *S1pr1*^high^ astrocytes by approximately 20% (**Fig. 3D**). Notably, this effect was not observed in the sample group treated with ozanimod alone or ozanimod with cisplatin, suggesting that S1PR1 is critical for the increased *S1pr1*^high^ subpopulation that may contribute to CIPN. We also found that the overall astrocytic *S1pr1* expression in the cisplatin-treated sample group was similar to that of the vehicle sample group, which may be attributed to the decreased *S1pr1* expression in the *Gfap*^high^ astrocyte subpopulation (**Fig. 3E**).

### *S1pr1*^high^ astrocytes-specific Wnt signaling contributes to CIPN in a S1PR1-dependent manner

Differential expression analysis of spinal astrocytes from ozanimod-pretreated mice in the CIPN model revealed 156 genes that were protected by ozanimod from cisplatin-induced upregulation (**Fig. 4A**). These genes were enriched in cell cycle G1/S phase transition, Wnt signaling, negative regulation of gliogenesis, Ras protein signaling and GPCR signaling pathways (**Fig. 4B**). Similar analysis revealed 118 genes that were protected by ozanimod from cisplatin-induced downregulation; the enriched pathways included cytoplasmic translation, protein folding, and cellular response to interleukin-4 and steroid hormones (**Fig. 4C,D**). The results indicated that cisplatin regulation of these genes and associated pathways depends on S1PR1 activity.

**Figure 4.**
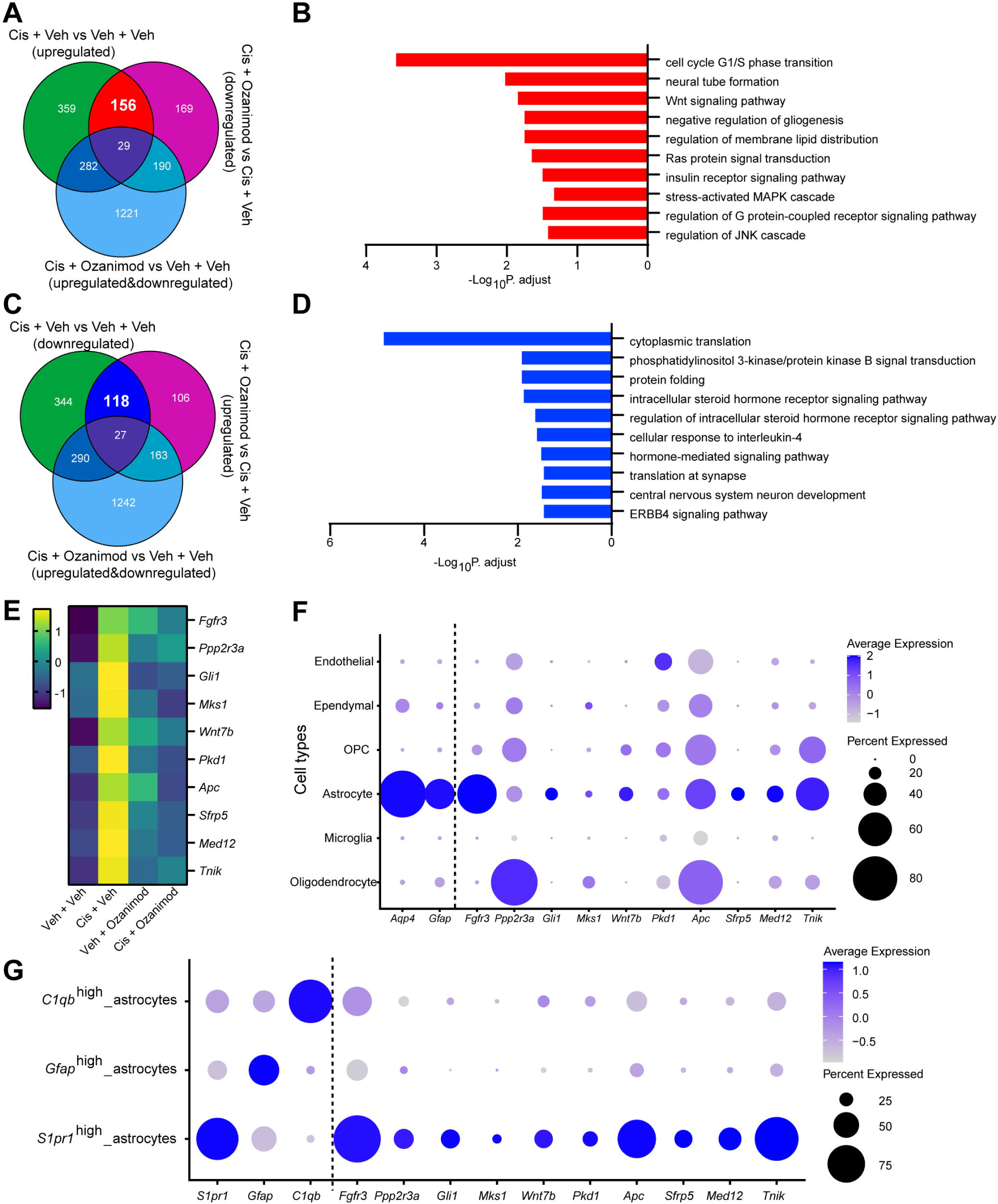
Ozanimod prevents the upregulation of astrocytic Wnt signaling from cisplatin. (A) Venn diagram showing 156 genes upregulated by cisplatin alone, but not with ozanimod pretreatment. (B) Enrichment pathway analysis of cisplatin-upregulated genes that were prevented by ozanimod, using cutoff adjusted p-value (P. adjust) < 0.05. (C) Venn diagram showing 118 genes that were downregulated by cisplatin alone, but not with ozanimod pretreatment. (D) Enrichment pathway analysis of cisplatin-downregulated genes that were prevented by ozanimod, using cutoff P. adjust < 0.05. (E) Heatmap of log2 normalized gene counts of Wnt signaling genes altered by cisplatin in an S1PR1-dependent manner. (F) Dotplot showing the expression profiles of Wnt signaling genes shown in (E) in all the identified cell types. Genes on the left side of dotted line are astrocyte-specific. (G) Dotplot showing the expression profiles of Wnt signaling genes shown in (E) in astrocyte subpopulations. Genes on the left side of dotted line are astrocyte subpopulation-specific.

The S1PR1-dependent cisplatin-upregulated Wnt signaling pathway genes included *Fgfr3*, *Ppp2r3a*, *Gli1*, *Mks1*, *Wnt7b*, *Pkd1*, *Apc*, *Sfrp5*, *Med12*, and *Tnik* (**Fig. 4E**). More than half of these genes showed astrocyte-specific expression patterns, including *Fgfr3*, *Gli1*, *Wnt7b*, *Sfrp5*, *Med12*, and *Tnik* (**Fig. 4F**). Notably, all of the 10 Wnt signaling genes were included in the 132 genes comprising the *S1pr1*^high^ astrocyte-specific Wnt signaling pathway described above. Consistently, all of them were highly expressed in the *S1pr1*^high^ astrocyte subpopulation (**Fig. 4G**).

To validate our scRNA seq data, RNAscope assays using *Fgfr3*, *S1pr1*, and *Gfap* probes were performed across all sample groups. Consistently, *Fgfr3* highly co-expresses with *S1pr1* among the sample groups (**Fig. 5A**). Proportion analysis of cells with different *S1pr1* expression levels in all *S1pr1*-positive cells revealed a cisplatin-induced increase in the population of *S1pr1*-positive cells with higher *S1pr1* expression levels (counts ≥ 20), which was prevented by ozanimod pretreatment (**Fig. 5B**), supporting the scRNA-seq data described above (**Fig. 3D**). We quantified *Fgfr3* mRNA signals in *S1pr1*-positive cells and *Gfap*-positive cells separately and found that *Fgfr3* was significantly induced by cisplatin in *S1pr1*-associated cells, but not in *Gfap*-associated cells (**Fig. 5C; Supplemental Fig. 4A,B**). Moreover, the induction of *Fgfr3* in *S1pr1*-associated astrocytes was prevented by ozanimod (**Fig. 5C**).

**Figure 5.**
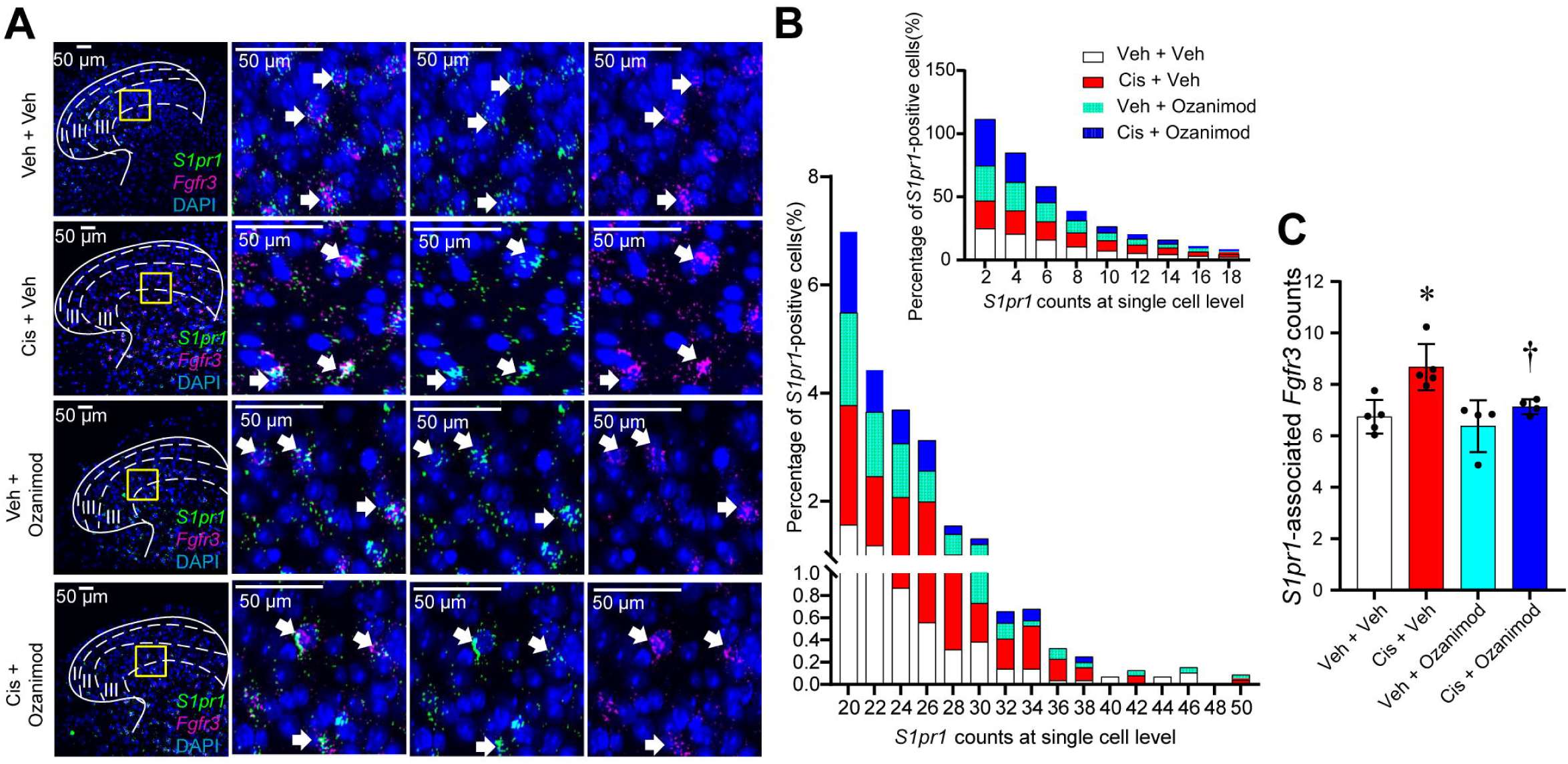
Ozanimod prevents the cisplatin-induced upregulation of *Fgfr3* gene expression in *S1pr1*-pisitive cells of mouse DH-SC. (A) Representative confocal images (left panel) showing expression of *S1pr1* (green) and *Fgfr3* (magenta) in DH-SC. DAPI indicates nuclear staining (blue). Scale bars 50 µm. Dotted lines represent boundaries of the first three dorsal horn laminae. Zoomed-in images of the yellow square area (100 µm × 100 µm) are shown in other panels. White arrows indicate the co-expression of *S1pr1* (green) and *Fgfr3* (magenta). (B) Percentage of cells with different levels of *S1pr1* expression levels in all *S1pr1* positive cells. Top right corner, *S1pr1* mRNA counts < 20; bottom left corner, *S1pr1* mRNA counts in a single cell level ≥ 20. (C) Quantification of *Fgfr3* mRNA signals in *S1pr1* positive cells of the I, II, and III laminae area of DH-SC. Data are mean ± SD for n = 4-5 mice/group; Ordinary one-way ANOVA with Tukey’s multiple comparison; *P < 0.05 vs. Veh + Veh, † P < 0.05 vs. Cis + Veh. Animal numbers of individual groups were indicated with the numbers of dots in the corresponding bars.

We performed immunofluorescence assays to examine the effect of cisplatin on FGFR3 and GFAP protein expression levels in DH-SC of CIPN mice. FGFR3 protein signals co-localized with GFAP protein signals at certain levels, even though they did not display GFAP-like intermediate filament structures [47] (**Fig. 6A,B**). A trend of increased FGFR3 protein levels by cisplatin was revealed, although it was not statistically significant **(Fig. 6C)**. However, FGFR3 protein levels were dramatically reduced in mice that received both ozanimod and cisplatin compared to mice treated with cisplatin only, which is consistent with the scRNA seq data described above **(Fig. 6C)**.

**Figure 6.**
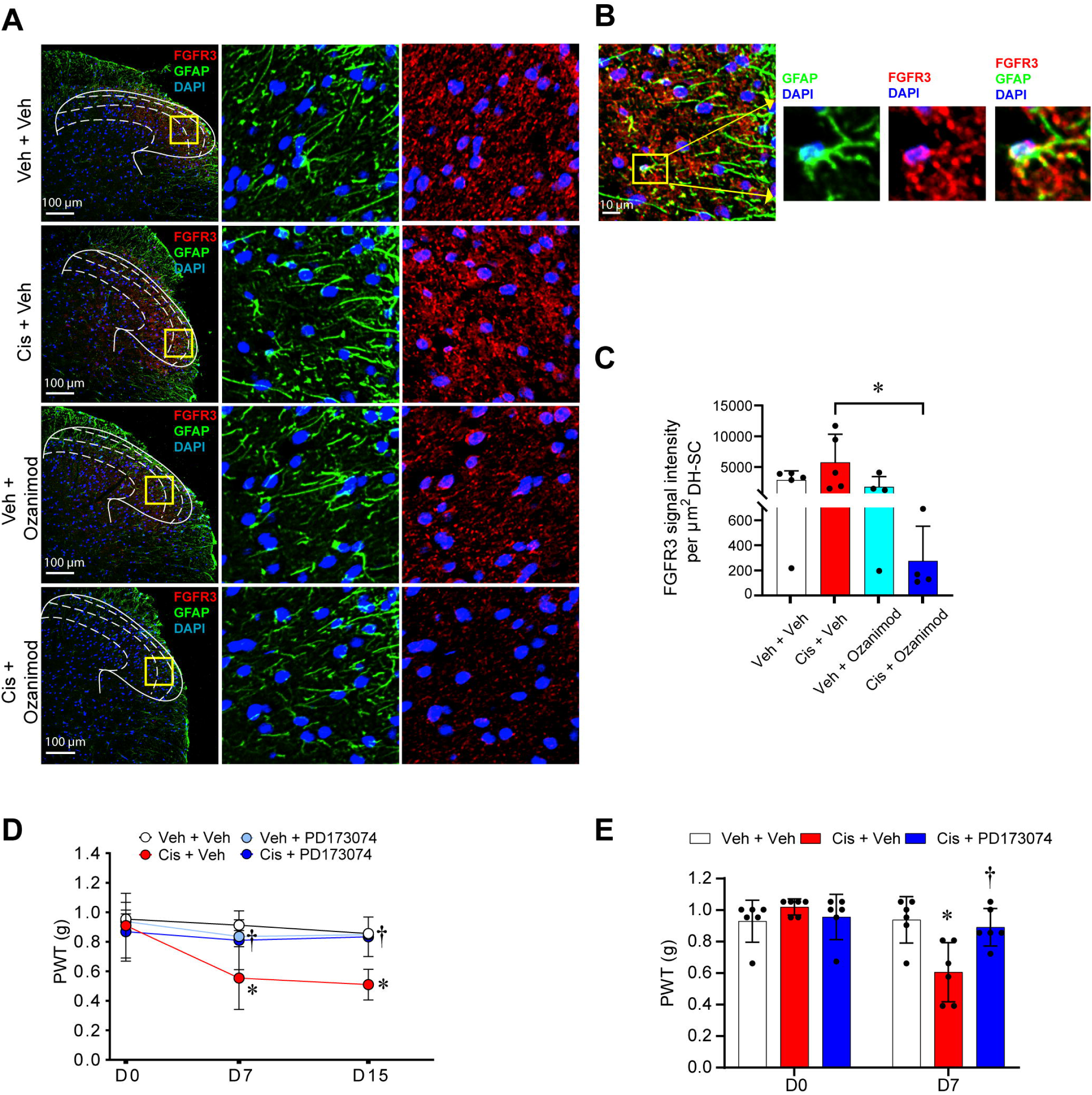
Cisplatin-induced neuropathic pain is prevented by FGFR3 antagonist, PD173074. (A) Representative confocal images (left panel) showing expression of GFAP (green) and FGFR3 (red) in DH-SC. DAPI indicates nuclear staining (blue). Scale bars 100 µm. Dotted lines represent boundaries of the first three dorsal horn laminae. Zoomed-in images of the yellow square area (100 µm × 100 µm) were shown in the middle (GFAP with DAPI) and right (FGFR3 with DAPI) panels. (B) Zoomed-in images of the yellow square area in the left panel showing the expression of GFAP (green) and FGFR3 (red) in astrocytes. Scale bar 10 µm. (C) Quantification of FGFR3 signal intensities in the I, II, and III laminae area of DH-SC. Data are mean ± SD for n = 4∼5 mice/group; Ordinary one-way ANOVA with Tukey’s multiple comparisons test; *P < 0.05. Animal numbers of individual groups were indicated with the numbers of dots in the corresponding bars. (D) Experimental paradigm for the treatment of PD173074 (i.p.) on CIPN mice. Male mice were treated with cisplatin or vehicle for two cycles of 5 daily doses of 2.3 mg/kg, i.p with 5 days of rest in between. PD173074 (20 mg/kg, i.p.) was administered 15 minutes before each cisplatin injection. Mechano-hypersensitivity was assessed before the start of the treatment (D0), 72h after the end of the first cycle of cisplatin (D7) and one day after the last treatment (D15). Data are mean ± SD for n = 6 mice/group; Two-way ANOVA with Tukey’s multiple comparisons test; *P < 0.05 versus Veh + Veh, †P < 0.05 versus Cis + Veh. (E) Experimental paradigm for the treatment of PD173074 (i.th.) on CINP mice. Male mice were treated with cisplatin or vehicle for 5 daily doses of 2.3 mg/kg, i.p. PD173074 (10µl 5µM, i.th.) administered 15 minutes before each cisplatin injection. Mechano-hypersensitivity was assessed before the start of the treatment (D0) and 72h after the end of the first cycle of cisplatin (D7). Data are mean ± SD for n = 6 mice/group; Two-way ANOVA with Tukey’s multiple comparisons test; *P < 0.05 versus Veh + Veh, †P < 0.05 versus Cis + Veh.

FGFR3 is a component of the FGF/FGFR signaling that often activates Wnt/β-catenin signaling to regulate various biological processes, such as cell proliferation and differentiation [41; 72]. However, the function of FGFR3 in the development of CIPN has not been elucidated. Intraperitoneal injections of the FGFR3 antagonist PD173074 to mice before each cisplatin treatment prevented cisplatin-induced mechano-allodynia (**Fig. 6D**). Moreover, intrathecal injections of PD173074 in mice during cisplatin treatment attenuated the development of allodynia identifying the spinal cord as a site of FGFR3 activity (**Fig. 6E**). Doses of PD173074 were chosen from previous studies [71; 76]. Overall, these results suggest a potential S1PR1/ FGFR3 link in the contribution of *S1pr1*^high^ astrocyte-specific Wnt signaling to CIPN.

### Ozanimod preserves the loss of intraepidermal nerve fibers (IENFs)

In addition to prominent effects of ozanimod observed at the level of the spinal cord, we also found important effects at the level of IENFs suggestive of a neuroprotective effect. Indeed, and consistent with previous reports [23; 35], we observed a significant loss in IENF density in the glabrous skin of the hind paw of mice treated with cisplatin when compared to vehicle using the pan-neuronal marker PGP9.5 [34] (**Fig. 7**). Ozanimod preserved the integrity of IENFs (**Fig. 7**).

**Figure 7.**
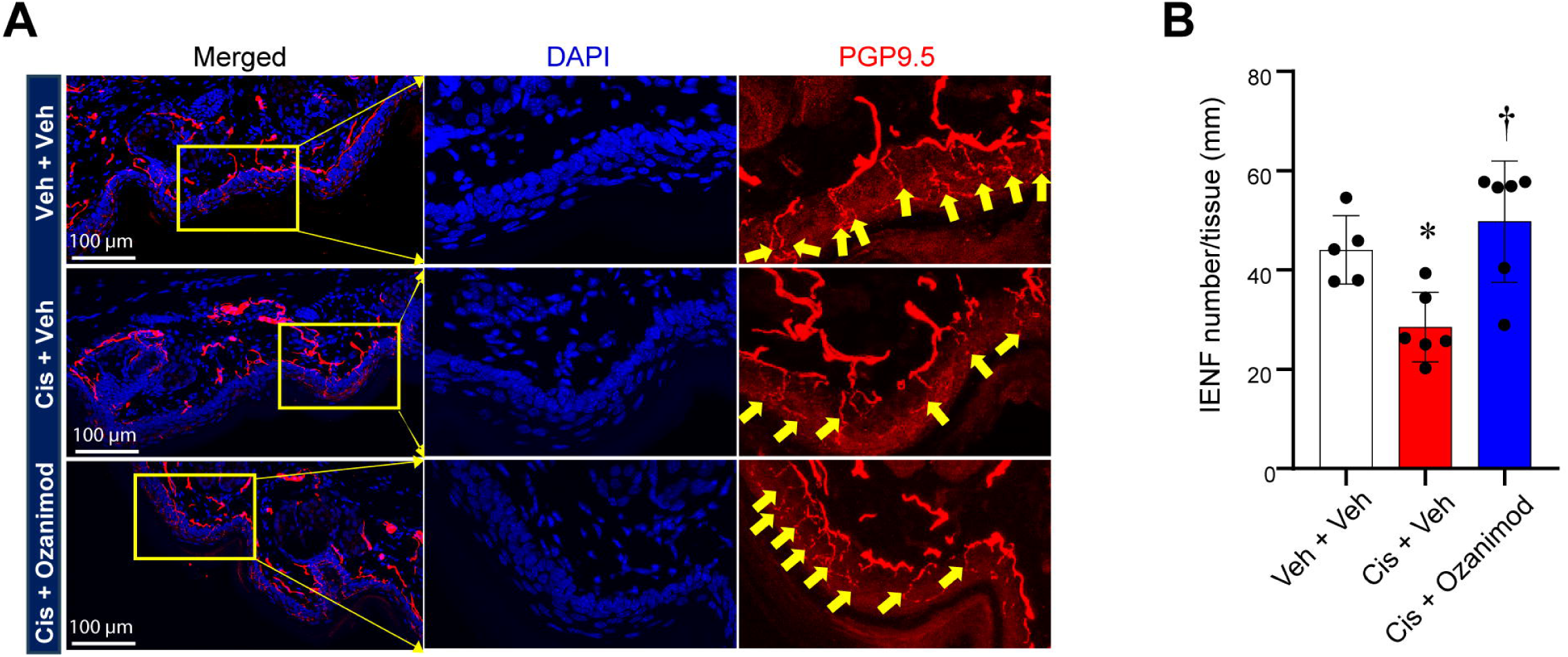
Preventive effect of ozanimod on cisplatin-induced IENF loss. (A) Representative confocal images showing immunofluorescence signals of PGP9.5 (red) and DAPI (blue) in the glabrous skin of the mouse hind paw. Scale bars 100 µm. Zoomed-in images were shown in the middle and right panel. IENFs are indicated with yellow arrows. (B) Quantitative data of IENFs density in different sample groups. IENFs density was calculated by dividing IENF number with length of the skin tissue. Mean±SD, n=5∼6/group, Ordinary one-way ANOVA with Tukey’s multiple comparisons test; *P<0.05 vs. Veh + Veh, † P<0.05 vs. Cis + Veh. Animal numbers of individual groups were indicated with the numbers of dots in the corresponding bars.

## Discussion

Activation of spinal astrocytes contributes to chemotherapy-induced peripheral neuropathy at least in part through promoting cytokine release [45; 51; 69]. Chemotherapeutic agents bortezomib and oxaliplatin failed to induce mechano-hypersensitivity when co-treated with glia activation inhibitor (minocycline) and astrocyte gap junctions decoupler (carbenoxolone) [50; 73]. Astrocyte activation is often associated with increased expression of GFAP protein in DH-SC. GFAP is the glial fibrillary acidic protein that provides mechanical support to astrocyte structure and the astroglial brain blood boundary (BBB). It mainly labels the extended branching of astrocytes and has been widely used as a protein marker for astrocytes [30; 38; 62]. Increased GFAP in tissues is also widely used to indicate cell proliferation and therefore reactivity [29].

We previously reported that expression of *S1pr1* in astrocytes is critical for the induction of neuropathic pain in several animal models of neuropathic pain including with chemotherapy (Chen et al., 2019; Stockstill et al., 2018). However, S1PR1 was rarely used as a cell marker for astrocytes due to the limited source of specific and reliable antibodies against GPCR proteins [36; 48].

Through single-cell RNA sequencing analysis, we found that neither *Gfap* nor *S1pr1* was highly expressed in all astrocytes. Instead, *Gfap* and *S1pr1* were highly expressed in different astrocyte subpopulations. Consistent with our scRNA-seq data, a recent study clustering astrocytes from mouse brain tissues identified *S1pr1* and *Vim*, a gene highly co-expressed with *Gfap* [42], as transcriptional signatures for two different astrocytes subpopulations [66]. We found that the *S1pr1*^high^ astrocyte subpopulation represented about half of the total astrocytes and the proportion was increased by cisplatin treatment. This effect did not lead to a significant increase in astrocytic *S1pr1* expression on average, which is not controversial given the decreased *S1pr1* expression in the *Gfap*^high^ astrocyte subpopulation. Increase in the *S1pr1*^high^ astrocyte subpopulation depended on S1PR1 signaling, since it was blocked by ozanimod. Increased *S1pr1*^high^ astrocyte subpopulations may be associated with an increased number of activated astrocytes, leading to persisting mechano-hypersensitivity.

Previous studies on downstream targets of S1PR1 signaling in CNS identified the p38-MAPK pathway, nuclear factor kappa-B, and the nod-like receptor family, pyrin domain containing 3 inflammasomes [12; 21; 60; 61]. However, these targets were normalized into all the spinal cells and could not be assigned to specific cell types. In this study, we demonstrated that a *S1pr1*^high^ astrocyte-specific Wnt signaling pathway, featuring *Wnt7b*, *Gli1*, *F*gfr3, and other seven genes that were highly expressed in the *S1pr1*^high^ astrocyte subpopulation, was upregulated by cisplatin in a S1PR1 signaling-dependent manner. Wnt signaling activates target genes through nuclear translocation of β-catenin [31]. It is triggered by the binding of ligands to cell surface receptors to regulate numerous biological processes such as cell cycle and migration [32]. Through this, Wnt signaling controls fundamental processes ranging from neurogenesis to BBB integrity in the CNS [1]. Wnt signaling has been implicated in neurodegenerative diseases, including Alzheimer’s disease [1], highlighting the importance of appropriate regulation of Wnt signaling. Wnt inhibition also causes analgesic effects in other neuropathic pain models including nerve ligation and chronic constriction injury [67; 75]. Our scRNA seq data suggested for the first time a possible connection between S1PR1 and Wnt signaling in the development of CIPN. Interestingly, *S1pr1* is a transcriptional target of β-catenin [44]. Thus, we propose that cisplatin treatment establishes a positive feedback loop between S1PR1 and Wnt signaling to promote *S1pr1* transcription at single cell levels, resulting in an increased proportion of *S1pr1*^high^ astrocytes, within which the upregulated Wnt signaling may contribute to CIPN persistence (**Fig. 8**).

**Figure 8.**
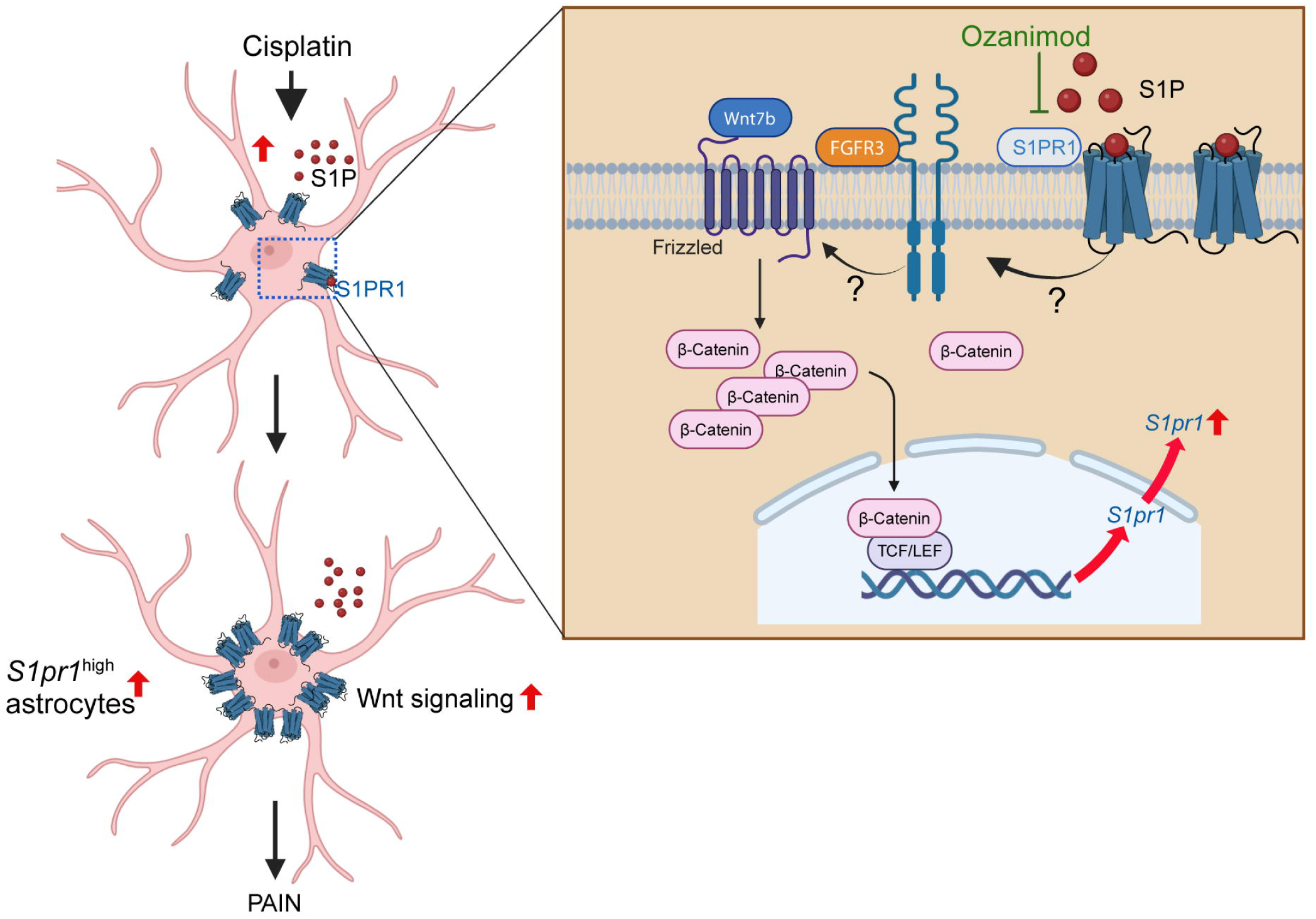
Schematic illustration showing contribution of *S1pr1*^high^ astrocytes to cisplatin-induced neuropathic pain. Cisplatin induces S1P release in DHSC. Binding of S1P to receptor (S1PR1) in astrocytes activates S1PR1 signaling, promoting FGFR3 and Wnt signaling, which increases the transcription of *S1pr1* through the nuclear translocation of β-Catenin. The resulting increase in proportion of *S1pr1*^high^ astrocytes and Wnt signaling provoke astrocyte activation, leading to neuropathic pain. Ozanimod abolishes this effect by inhibiting S1PR1signaling. Figure was created with Biorender.com

FGFR3 is one of the FGF/FGFR family members that regulate Wnt/β-catenin signaling in various biological processes [14; 17; 22]. In addition to the identical expression profile of *Fgfr3* to *S1pr1* in astrocyte subpopulations, the S1PR1-dependent upregulation of *Fgfr3* by cisplatin suggests FGFR3 as a potential S1PR1 signaling target. The absence of mechano-hypersensitivity in mice treated with cisplatin and the FGFR3 antagonist, PD170734, further supports our hypothesis. Consistently, studies in the spared nerve injury model of neuropathic pain revealed that spinal astrocytic FGFR3 activation leads to mechano-hypersensitivity through tumor necrosis factor alpha [71]. Therefore, we propose that FGFR3 plays a bridging role connecting S1PR1 and Wnt signaling in response to cisplatin (**Fig. 8**). However, it is an open question if FGFR3 directly activates or works in parallel with Wnt signaling to induce CIPN. Thus, future studies will investigate the mechanism underlying the S1PR1-mediated upregulation of *Fgfr3* and Wnt signaling. In addition to effects on spinal cord astrocytes, ozanimod also prevented the loss in IENF, suggesting neuroprotective effects. The mechanisms involved remain to be investigated but may include protective effects on the mitochondria or inhibition of Wnt signaling. For example, S1PR1 antagonists have been shown in previous studies including ours to have mitoprotective effects [60] and Wnt inhibitors attenuate CIPN and significantly improve chemotherapy-induced reduction in IENF density [49]. Overall, our findings bridge gaps in our understanding of cell and molecular changes induced by cisplatin, characterized astrocytes heterogeneity, and identified a new target for therapeutic intervention addressing a pressing medical need. On a broader level, our work is anticipated to drive the investigation of these pathways in other cancer treatment-associated neurotoxicities such as cognitive deficits where an astrocyte based S1P/S1PR1 axis has been implicated [60].

## Acknowledgements

We thank Drs. Eissenberg and Doyle (Saint Louis University School of Medicine) for their input and review of this manuscript. This study was funded by SLU startup funds (to DS) and NIH grant R01CA261979 (DS). None of the authors have a conflict of interest.

## References

[1] Aghaizu ND, Jin H, Whiting PJ. Dysregulated Wnt Signalling in the Alzheimer’s Brain. Brain Sci 2020;10(12).

[2] Bencardino S, D’Amico F, Faggiani I, Bernardi F, Allocca M, Furfaro F, Parigi TL, Zilli A, Fiorino G, Peyrin-Biroulet L, Danese S. Efficacy and Safety of S1P1 Receptor Modulator Drugs for Patients with Moderate-to-Severe Ulcerative Colitis. J Clin Med 2023;12(15).

[3] Bennett GJ, Doyle T, Salvemini D. Mitotoxicity in distal symmetrical sensory peripheral neuropathies. Nat Rev Neurol 2014;10(6):326–336.

[4] Brandolini L, d’Angelo M, Antonosante A, Allegretti M, Cimini A. Chemokine Signaling in Chemotherapy-Induced Neuropathic Pain. Int J Mol Sci 2019;20(12).

[5] Brown A, Kumar S, Tchounwou PB. Cisplatin-Based Chemotherapy of Human Cancers. J Cancer Sci Ther 2019;11(4).

[6] Cavaletti G, Marmiroli P. Chemotherapy-induced peripheral neurotoxicity. Curr Opin Neurol 2015;28(5):500–507.

[7] Chaplan SR, Bach FW, Pogrel JW, Chung JM, Yaksh TL. Quantitative assessment of tactile allodynia in the rat paw. J Neurosci Methods 1994;53(1):55–63.

[8] Chen Z, Doyle TM, Luongo L, Largent-Milnes TM, Giancotti LA, Kolar G, Squillace S, Boccella S, Walker JK, Pendleton A, Spiegel S, Neumann WL, Vanderah TW, Salvemini D. Sphingosine-1-phosphate receptor 1 activation in astrocytes contributes to neuropathic pain. Proc Natl Acad Sci U S A 2019;116(21):10557–10562.

[9] Chiu GS, Maj MA, Rizvi S, Dantzer R, Vichaya EG, Laumet G, Kavelaars A, Heijnen CJ. Pifithrin-mu Prevents Cisplatin-Induced Chemobrain by Preserving Neuronal Mitochondrial Function. Cancer Res 2017;77(3):742–752.

[10] Colvin LA. Chemotherapy-induced peripheral neuropathy: where are we now? Pain 2019;160 Suppl 1(Suppl 1):S1–S10.

[11] Dixon WJ. Efficient analysis of experimental observations. Annu Rev Pharmacol Toxicol 1980;20:441–462.

[12] Doyle TM, Chen Z, Durante M, Salvemini D. Activation of Sphingosine-1-Phosphate Receptor 1 in the Spinal Cord Produces Mechanohypersensitivity Through the Activation of Inflammasome and IL-1beta Pathway. J Pain 2019;20(8):956–964.

[13] Doyle TM, Salvemini D. Mini-Review: Mitochondrial dysfunction and chemotherapy-induced neuropathic pain. Neurosci Lett 2021;760:136087.

[14] Goto H, Kimmey SC, Row RH, Matus DQ, Martin BL. FGF and canonical Wnt signaling cooperate to induce paraxial mesoderm from tailbud neuromesodermal progenitors through regulation of a two-step epithelial to mesenchymal transition. Development 2017;144(8):1412–1424.

[15] Grace PM, Hutchinson MR, Maier SF, Watkins LR. Pathological pain and the neuroimmune interface. Nat Rev Immunol 2014;14(4):217–231.

[16] Gutbier S, Spreng AS, Delp J, Schildknecht S, Karreman C, Suciu I, Brunner T, Groettrup M, Leist M. Prevention of neuronal apoptosis by astrocytes through thiol-mediated stress response modulation and accelerated recovery from proteotoxic stress. Cell Death Differ 2018;25(12):2101–2117.

[17] Harshuk-Shabso S, Dressler H, Niehrs C, Aamar E, Enshell-Seijffers D. Fgf and Wnt signaling interaction in the mesenchymal niche regulates the murine hair cycle clock. Nat Commun 2020;11(1):5114.

[18] Inoue K, Tsuda M. Microglia in neuropathic pain: cellular and molecular mechanisms and therapeutic potential. Nat Rev Neurosci 2018;19(3):138–152.

[19] Janes K, Little JW, Li C, Bryant L, Chen C, Chen Z, Kamocki K, Doyle T, Snider A, Esposito E, Cuzzocrea S, Bieberich E, Obeid L, Petrache I, Nicol G, Neumann WL, Salvemini D. The development and maintenance of paclitaxel-induced neuropathic pain require activation of the sphingosine 1-phosphate receptor subtype 1. J Biol Chem 2014;289(30):21082–21097.

[20] Koizumi S, Fujishita K, Inoue K. Regulation of cell-to-cell communication mediated by astrocytic ATP in the CNS. Purinergic Signal 2005;1(3):211–217.

[21] Komiya H, Takeuchi H, Ogasawara A, Ogawa Y, Kubota S, Hashiguchi S, Takahashi K, Kunii M, Tanaka K, Tada M, Doi H, Tanaka F. Siponimod inhibits microglial inflammasome activation. Neurosci Res 2025.

[22] Krejci P, Aklian A, Kaucka M, Sevcikova E, Prochazkova J, Masek JK, Mikolka P, Pospisilova T, Spoustova T, Weis M, Paznekas WA, Wolf JH, Gutkind JS, Wilcox WR, Kozubik A, Jabs EW, Bryja V, Salazar L, Vesela I, Balek L. Receptor tyrosine kinases activate canonical WNT/beta-catenin signaling via MAP kinase/LRP6 pathway and direct beta-catenin phosphorylation. PLoS One 2012;7(4):e35826.

[23] Krukowski K, Ma J, Golonzhka O, Laumet GO, Gutti T, van Duzer JH, Mazitschek R, Jarpe MB, Heijnen CJ, Kavelaars A. HDAC6 inhibition effectively reverses chemotherapy-induced peripheral neuropathy. Pain 2017;158(6):1126–1137.

[24] Kunkel GT, Maceyka M, Milstien S, Spiegel S. Targeting the sphingosine-1-phosphate axis in cancer, inflammation and beyond. Nat Rev Drug Discov 2013;12(9):688–702.

[25] Lamb YN. Ozanimod: First Approval. Drugs 2020;80(8):841–848.

[26] Laumet G, Bavencoffe A, Edralin JD, Huo XJ, Walters ET, Dantzer R, Heijnen CJ, Kavelaars A. Interleukin-10 resolves pain hypersensitivity induced by cisplatin by reversing sensory neuron hyperexcitability. Pain 2020;161(10):2344–2352.

[27] Lees JG, Makker PG, Tonkin RS, Abdulla M, Park SB, Goldstein D, Moalem-Taylor G. Immune-mediated processes implicated in chemotherapy-induced peripheral neuropathy. Eur J Cancer 2017;73:22–29.

[28] Li C, Wu Z, Zhou L, Shao J, Hu X, Xu W, Ren Y, Zhu X, Ge W, Zhang K, Liu J, Huang R, Yu J, Luo D, Yang X, Zhu W, Zhu R, Zheng C, Sun YE, Cheng L. Temporal and spatial cellular and molecular pathological alterations with single-cell resolution in the adult spinal cord after injury. Signal Transduct Target Ther 2022;7(1):65.

[29] Liddelow SA, Barres BA. Reactive Astrocytes: Production, Function, and Therapeutic Potential. Immunity 2017;46(6):957–967.

[30] Liedtke W, Edelmann W, Bieri PL, Chiu FC, Cowan NJ, Kucherlapati R, Raine CS. GFAP is necessary for the integrity of CNS white matter architecture and long-term maintenance of myelination. Neuron 1996;17(4):607–615.

[31] Liu J, Xiao Q, Xiao J, Niu C, Li Y, Zhang X, Zhou Z, Shu G, Yin G. Wnt/beta-catenin signalling: function, biological mechanisms, and therapeutic opportunities. Signal Transduct Target Ther 2022;7(1):3.

[32] Logan CY, Nusse R. The Wnt signaling pathway in development and disease. Annu Rev Cell Dev Biol 2004;20:781–810.

[33] Maceyka M, Harikumar KB, Milstien S, Spiegel S. Sphingosine-1-phosphate signaling and its role in disease. Trends Cell Biol 2012;22(1):50–60.

[34] Maj MA, Ma J, Krukowski KN, Kavelaars A, Heijnen CJ. Inhibition of Mitochondrial p53 Accumulation by PFT-mu Prevents Cisplatin-Induced Peripheral Neuropathy. Front Mol Neurosci 2017;10:108.

[35] Mao-Ying QL, Kavelaars A, Krukowski K, Huo XJ, Zhou W, Price TJ, Cleeland C, Heijnen CJ. The anti-diabetic drug metformin protects against chemotherapy-induced peripheral neuropathy in a mouse model. PLoS One 2014;9(6):e100701.

[36] Michel MC, Wieland T, Tsujimoto G. How reliable are G-protein-coupled receptor antibodies? Naunyn Schmiedebergs Arch Pharmacol 2009;379(4):385–388.

[37] Milich LM, Choi JS, Ryan C, Cerqueira SR, Benavides S, Yahn SL, Tsoulfas P, Lee JK. Single-cell analysis of the cellular heterogeneity and interactions in the injured mouse spinal cord. J Exp Med 2021;218(8).

[38] Miller RH, Raff MC. Fibrous and protoplasmic astrocytes are biochemically and developmentally distinct. J Neurosci 1984;4(2):585–592.

[39] Morrens J, Van Den Broeck W, Kempermann G. Glial cells in adult neurogenesis. Glia 2012;60(2):159–174.

[40] Newton J, Lima S, Maceyka M, Spiegel S. Revisiting the sphingolipid rheostat: Evolving concepts in cancer therapy. Exp Cell Res 2015;333(2):195–200.

[41] Nguyen TM, Kabotyanski EB, Dou Y, Reineke LC, Zhang P, Zhang XH, Malovannaya A, Jung SY, Mo Q, Roarty KP, Chen Y, Zhang B, Neilson JR, Lloyd RE, Perou CM, Ellis MJ, Rosen JM. FGFR1-Activated Translation of WNT Pathway Components with Structured 5’ UTRs Is Vulnerable to Inhibition of EIF4A-Dependent Translation Initiation. Cancer Res 2018;78(15):4229–4240.

[42] O’Leary LA, Davoli MA, Belliveau C, Tanti A, Ma JC, Farmer WT, Turecki G, Murai KK, Mechawar N. Characterization of Vimentin-Immunoreactive Astrocytes in the Human Brain. Front Neuroanat 2020;14:31.

[43] Old EA, Malcangio M. Chemokine mediated neuron-glia communication and aberrant signalling in neuropathic pain states. Curr Opin Pharmacol 2012;12(1):67–73.

[44] Oliveira-Paula GH, Liu S, Maira A, Ressa G, Ferreira GC, Quintar A, Jayakumar S, Almonte V, Parikh D, Valenta T, Basler K, Hla T, Riascos-Bernal DF, Sibinga NES. The beta-catenin C terminus links Wnt and sphingosine-1-phosphate signaling pathways to promote vascular remodeling and atherosclerosis. Sci Adv 2024;10(11):eadg9278.

[45] Park HJ, Stokes JA, Corr M, Yaksh TL. Toll-like receptor signaling regulates cisplatin-induced mechanical allodynia in mice. Cancer Chemother Pharmacol 2014;73(1):25–34.

[46] Park SB, Goldstein D, Krishnan AV, Lin CS, Friedlander ML, Cassidy J, Koltzenburg M, Kiernan MC. Chemotherapy-induced peripheral neurotoxicity: a critical analysis. CA Cancer J Clin 2013;63(6):419–437.

[47] Pekny M, Pekna M. Astrocyte intermediate filaments in CNS pathologies and regeneration. J Pathol 2004;204(4):428–437.

[48] Pradidarcheep W, Stallen J, Labruyere WT, Dabhoiwala NF, Michel MC, Lamers WH. Lack of specificity of commercially available antisera against muscarinergic and adrenergic receptors. Naunyn Schmiedebergs Arch Pharmacol 2009;379(4):397–402.

[49] Resham K, Sharma SS. Pharmacological interventions targeting Wnt/beta-catenin signaling pathway attenuate paclitaxel-induced peripheral neuropathy. Eur J Pharmacol 2019;864:172714.

[50] Robinson CR, Dougherty PM. Spinal astrocyte gap junction and glutamate transporter expression contributes to a rat model of bortezomib-induced peripheral neuropathy. Neuroscience 2015;285:1–10.

[51] Robinson CR, Zhang H, Dougherty PM. Astrocytes, but not microglia, are activated in oxaliplatin and bortezomib-induced peripheral neuropathy in the rat. Neuroscience 2014;274:308–317.

[52] Rostami N, Nikkhoo A, Ajjoolabady A, Azizi G, Hojjat-Farsangi M, Ghalamfarsa G, Yousefi B, Yousefi M, Jadidi-Niaragh F. S1PR1 as a Novel Promising Therapeutic Target in Cancer Therapy. Mol Diagn Ther 2019;23(4):467–487.

[53] Ruggieri S, Quartuccio ME, Prosperini L. Ponesimod in the Treatment of Relapsing Forms of Multiple Sclerosis: An Update on the Emerging Clinical Data. Degener Neurol Neuromuscul Dis 2022;12:61–73.

[54] Santos EN, Fields RD. Regulation of myelination by microglia. Sci Adv 2021;7(50):eabk1131.

[55] Secci ME, Reed T, Quinlan V, Gilpin NW, Avegno EM. Quantitative Analysis of Gene Expression in RNAscope-processed Brain Tissue. Bio Protoc 2023;13(1):e4580.

[56] Sharma S, Mathur AG, Pradhan S, Singh DB, Gupta S. Fingolimod (FTY720): First approved oral therapy for multiple sclerosis. J Pharmacol Pharmacother 2011;2(1):49–51.

[57] Singh SK, Spiegel S. Sphingosine-1-phosphate signaling: A novel target for simultaneous adjuvant treatment of triple negative breast cancer and chemotherapy-induced neuropathic pain. Adv Biol Regul 2020;75:100670.

[58] Sprauten M, Darrah TH, Peterson DR, Campbell ME, Hannigan RE, Cvancarova M, Beard C, Haugnes HS, Fossa SD, Oldenburg J, Travis LB. Impact of long-term serum platinum concentrations on neuro– and ototoxicity in Cisplatin-treated survivors of testicular cancer. J Clin Oncol 2012;30(3):300–307.

[59] Spurgat MS, Tang SJ. Single-Cell RNA-Sequencing: Astrocyte and Microglial Heterogeneity in Health and Disease. Cells 2022;11(13).

[60] Squillace S, Niehoff ML, Doyle TM, Green M, Esposito E, Cuzzocrea S, Arnatt CK, Spiegel S, Farr SA, Salvemini D. Sphingosine-1-phosphate receptor 1 activation in the central nervous system drives cisplatin-induced cognitive impairment. J Clin Invest 2022;132(17).

[61] Squillace S, Spiegel S, Salvemini D. Targeting the Sphingosine-1-Phosphate Axis for Developing Non-narcotic Pain Therapeutics. Trends Pharmacol Sci 2020;41(11):851–867.

[62] Stavale LM, Soares ES, Mendonca MC, Irazusta SP, da Cruz Hofling MA. Temporal relationship between aquaporin-4 and glial fibrillary acidic protein in cerebellum of neonate and adult rats administered a BBB disrupting spider venom. Toxicon 2013;66:37–46.

[63] Stockstill K, Doyle TM, Yan X, Chen Z, Janes K, Little JW, Braden K, Lauro F, Giancotti LA, Harada CM, Yadav R, Xiao WH, Lionberger JM, Neumann WL, Bennett GJ, Weng HR, Spiegel S, Salvemini D. Dysregulation of sphingolipid metabolism contributes to bortezomib-induced neuropathic pain. J Exp Med 2018;215(5):1301–1313.

[64] Stockstill K, Wahlman C, Braden K, Chen Z, Yosten GL, Tosh DK, Jacobson KA, Doyle TM, Samson WK, Salvemini D. Sexually dimorphic therapeutic response in bortezomib-induced neuropathic pain reveals altered pain physiology in female rodents. Pain 2020;161(1):177–184.

[65] Sun J, Nan G. The extracellular signal-regulated kinase 1/2 pathway in neurological diseases: A potential therapeutic target (Review). Int J Mol Med 2017;39(6):1338–1346.

[66] Sun W, Liu Z, Jiang X, Chen MB, Dong H, Liu J, Sudhof TC, Quake SR. Spatial transcriptomics reveal neuron-astrocyte synergy in long-term memory. Nature 2024;627(8003):374–381.

[67] Tang J, Ji Q, Jin L, Tian M, Zhang LD, Liu XY. Secreted frizzled-related protein 1 regulates the progression of neuropathic pain in mice following spinal nerve ligation. J Cell Physiol 2018;233(8):5815–5822.

[68] Vendrell I, Macedo D, Alho I, Dionisio MR, Costa L. Treatment of Cancer Pain by Targeting Cytokines. Mediators Inflamm 2015;2015:984570.

[69] Wahlman C, Doyle TM, Little JW, Luongo L, Janes K, Chen Z, Esposito E, Tosh DK, Cuzzocrea S, Jacobson KA, Salvemini D. Chemotherapy-induced pain is promoted by enhanced spinal adenosine kinase levels through astrocyte-dependent mechanisms. Pain 2018;159(6):1025–1034.

[70] Wang XM, Lehky TJ, Brell JM, Dorsey SG. Discovering cytokines as targets for chemotherapy-induced painful peripheral neuropathy. Cytokine 2012;59(1):3–9.

[71] Xie KY, Wang Q, Cao DJ, Liu J, Xie XF. Spinal astrocytic FGFR3 activation leads to mechanical hypersensitivity by increased TNF-alpha in spared nerve injury. Int J Clin Exp Pathol 2019;12(8):2898–2908.

[72] Yin Y, White AC, Huh SH, Hilton MJ, Kanazawa H, Long F, Ornitz DM. An FGF-WNT gene regulatory network controls lung mesenchyme development. Dev Biol 2008;319(2):426–436.

[73] Yoon SY, Robinson CR, Zhang H, Dougherty PM. Spinal astrocyte gap junctions contribute to oxaliplatin-induced mechanical hypersensitivity. J Pain 2013;14(2):205–214.

[74] Yu G, Wang LG, Han Y, He QY. clusterProfiler: an R package for comparing biological themes among gene clusters. OMICS 2012;16(5):284–287.

[75] Zhao Y, Yang Z. Effect of Wnt signaling pathway on pathogenesis and intervention of neuropathic pain. Exp Ther Med 2018;16(4):3082–3088.

[76] Zheng Y, Ma H, Hu E, Huang Z, Cheng X, Xiong C. Inhibition of FGFR Signaling With PD173074 Ameliorates Monocrotaline-induced Pulmonary Arterial Hypertension and Rescues BMPR-II Expression. J Cardiovasc Pharmacol 2015;66(5):504–514.

