## Supplemental Figures for "Contribution of *S1pr1*-featured astrocyte subpopulation to cisplatin-induced neuropathic pain"

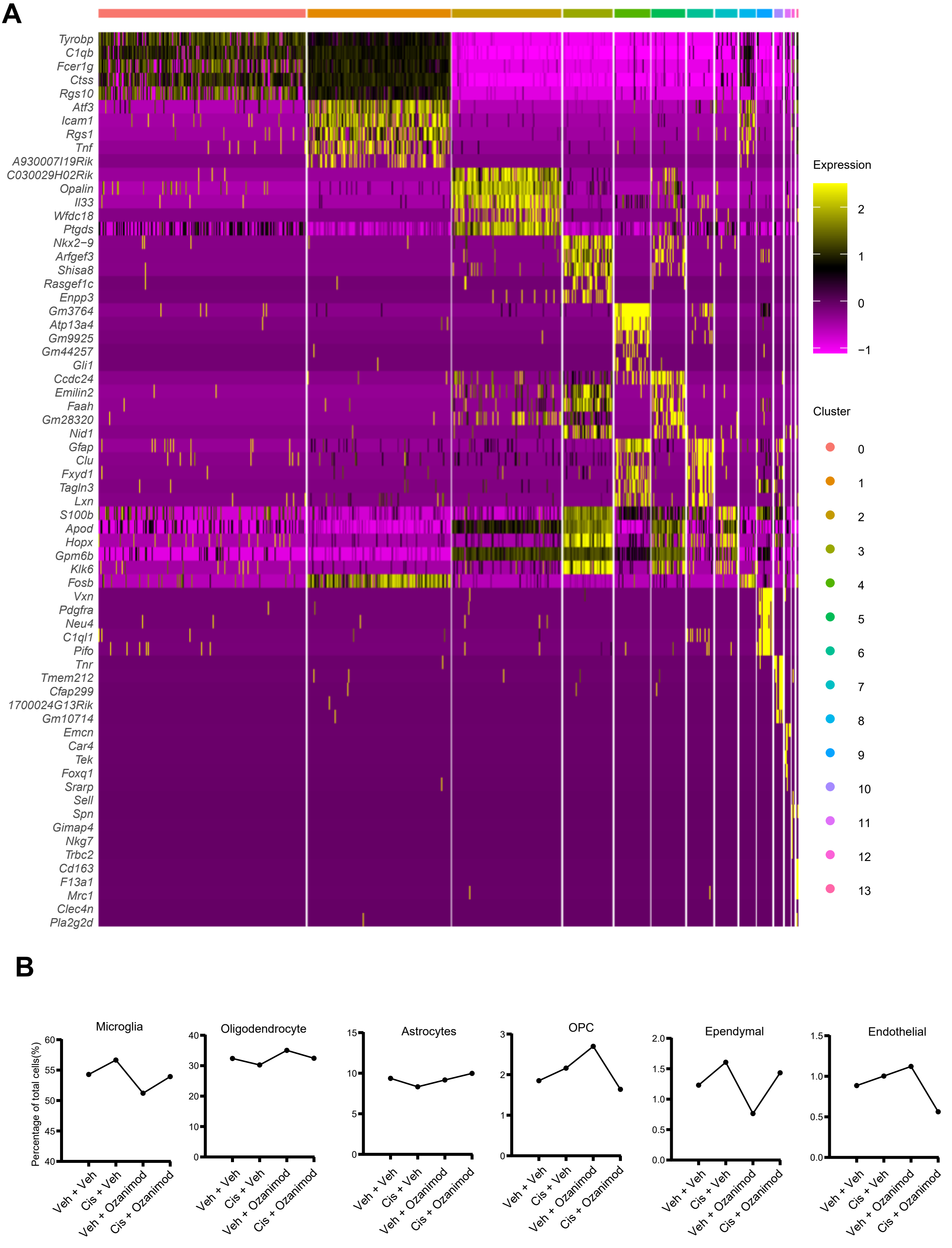


**Supplemental Figure 1.**
(A) Heatmap showing the expression of top 5 distinct markers for each cluster of spinal cells.
(B) Percentage of each cell type in total spinal cells across sample groups.

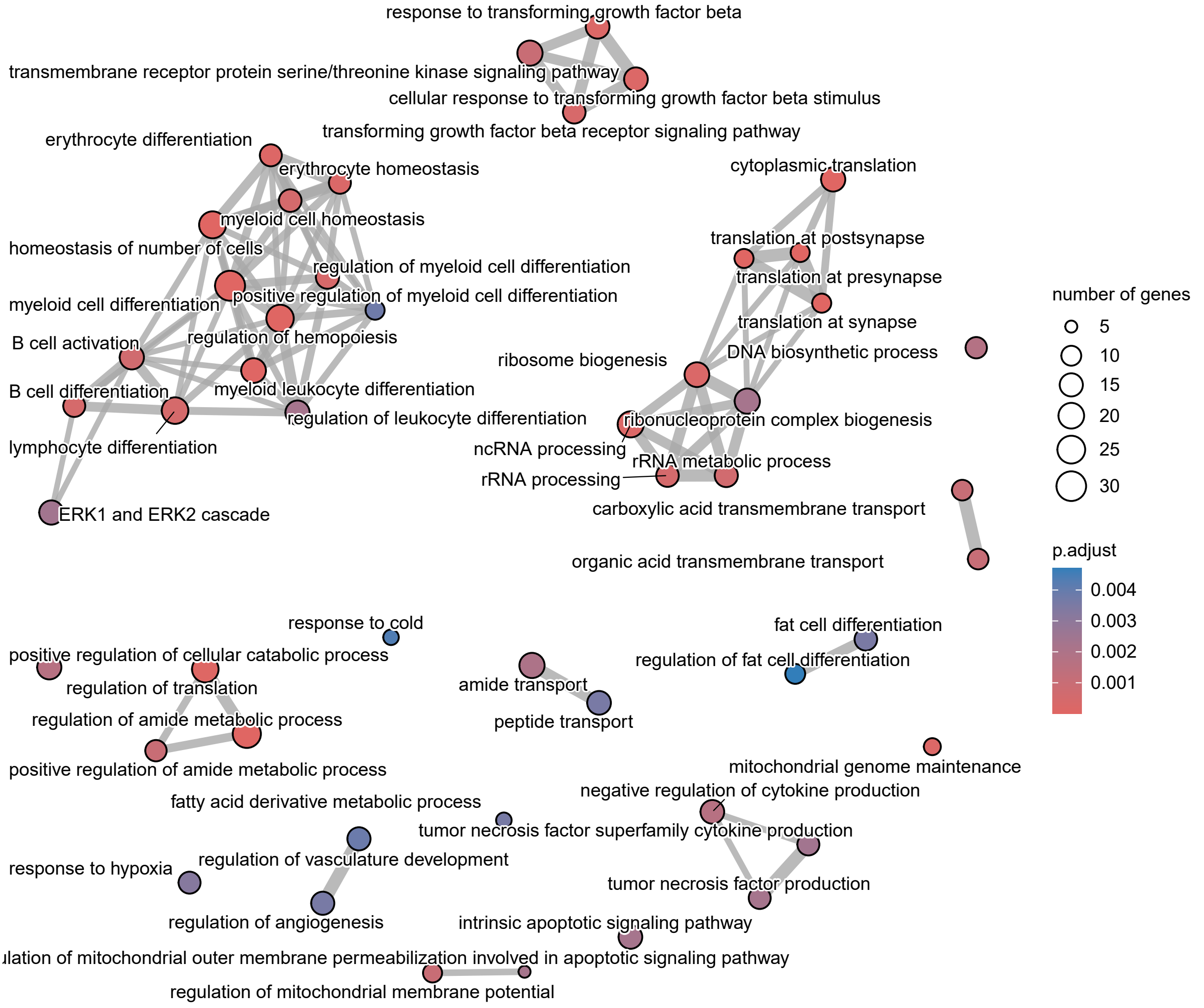


**Supplemental Figure 2.**
Enrichment map visualizing networks among cisplatin-upregulated pathways in microglia. Pathways are connected when there are overlaps in associated DEGs.

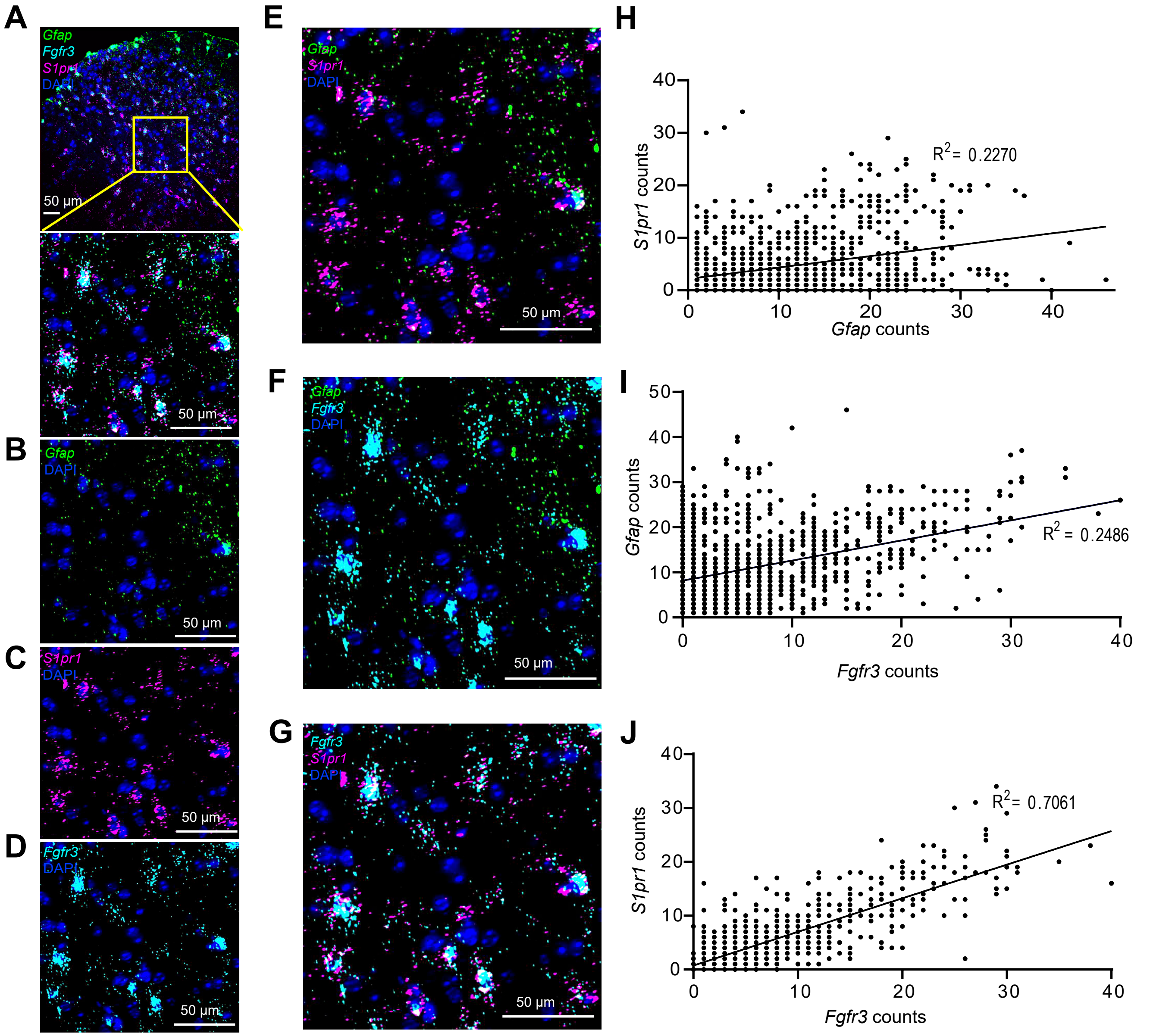


**Supplemental Figure 3. Distinct expression profiles of *S1pr1* and *Gfap* in mouse DH-SC.** (A) Top panel, representative confocal image of *S1pr1, Fgfr3,* and *Gfap* in mouse DH-SC. Bottom panel, zoomed-in image of the yellow square area in the top panel. *Gfap, S1pr1, Fgfr3* mRNA signals were shown with DAPI separately in (B-D). (E-G) Coexpression between different combinations of *S1pr1, Fgfr3,* and *Gfap* from the zoomed-in image in (A). (H-J) Liner regression and R square value showing different levels of proportional relationship between indicated mRNA signals at single cell level. Scale bar = 50 µm.

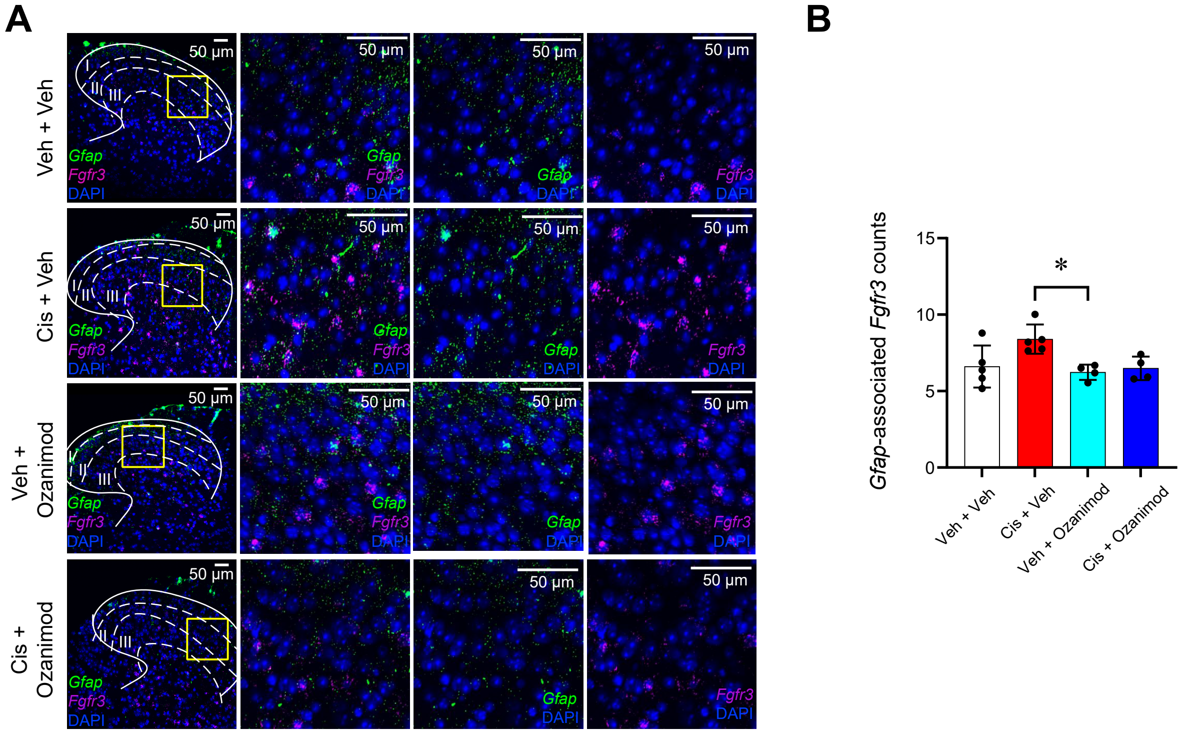


**Supplemental Figure 4. Visualization of gene expressions in mouse DH-SC.**
(A) Representative confocal images (left panel) showing expression of *Gfap* (green) and *Fgfr3* (magenta) in DH-SC. DAPI indicates nuclear staining (blue). Scale bars = 50 μm. Dotted lines represent boundaries of the first three dorsal horn laminae. Zoomed-in images of the yellow square area (150 μm × 150 μm) were showed in other panels. (B) Quantification of *Fgfr3* mRNA signals in *Gfap*-positive cells of I, II, and III laminae area. Data are mean ±SD; Ordinary one-way ANOVA with Tukey's multiple comparison; *$P < 0.05$ for n = 4~5 mice/group.
